# Human IGH germline gene diversity and allele frequencies in 2486 individuals from 25 global populations delineated by ultra-high throughput genotyping

**DOI:** 10.1101/2025.08.06.668935

**Authors:** Martin Corcoran, Sanjana Narang, Mateusz Kaduk, Mark Chernyshev, Anna Färnert, Christopher Sundling, Gunilla B. Karlsson Hedestam

## Abstract

The extraordinary diversity of human immunoglobulin (IG) genes underpins effective antibody responses, yet the full scope and functional impact of germline-encoded IG variation across populations is largely unknown. Here, we present ImmuneDiscover, an ultra-high-throughput sequencing platform enabling individualized IG genotyping from nanogram-scale DNA and simultaneous analysis of over 1,000 individuals. Using ImmuneDiscover, we generated high-resolution IG genotypes for 2,486 individuals from the 1000 Genomes Project, spanning diverse ancestries, and built KIARVA, an open-access atlas of IG gene variation. Cross-validation with SNP data from over one million genomes confirmed the accuracy and novelty of discovered alleles. Our analysis reveals striking population-specific patterns, including a multigene IGHD segment deletion found in up to 30% of East Asian individuals, and haplotypic differences linked to disease-associated loci. These findings illuminate the evolutionary forces shaping IG diversity and provide a foundational resource for investigating the immunogenetic basis of pathogen susceptibility and vaccine responsiveness worldwide.

## Introduction

The human adaptive immune system plays an essential role in countering infections and providing immunological memory, a major component of which involves B cells ^1^. The B cell receptor (BCR) genes, along with those encoding T cell receptors, are unique amongst human genes in that they exist in a state of non-functional, highly polymorphic germline-encoded *variable* (V), *diversity* (D) and *junctional* (J) variants that necessitate a process of genomic recombination before functional receptors can be expressed. In addition to the combinatorial rearrangement of these genes, nucleotides may be trimmed or added at the VDJ and VJ junctions, resulting in highly diverse repertoires of receptors in each person ^2^.

The immunoglobulin heavy chain (IGH) locus is exceptionally heterogeneous and has evolved through the acquisition of polymorphisms, gene duplications, deletions, and pseudogene formation ^3,4^. Specifically, the IG loci have been shaped by endemic and pandemic pathogens, and the concomitant ability to avoid recognition of self-antigens. The diverse range of binding capacities present in the naïve BCR repertoire can be further refined through one or more rounds of affinity maturation ^5^. This results in a delicate immunological calculus between BCRs that enable rapid responses to endemic pathogens and those that develop sufficient breadth through affinity maturation. While a given individual will have a greater or lesser ability to cope with a specific pathogen, the overall population has evolved to withstand exposures to multiple pathogens. This form of immunological heterogeneity is evident in the variety of MHC polymorphisms found both within and between diverse human populations ^6,7^. Two main public reference databases collating IG allelic variants are the international ImMunoGeneTics (IMGT) information system ^8,9^ and the Adaptive Immune Receptor Repertoire Community (AIRR-C) IG reference sets ^10^, both of which are heavily biased towards alleles of European origin. Therefore, a central unmet need is to define the range of IG germline variation in diverse human population groups ^11^. Leveraging this knowledge, in combination with accurate information on pathogen associated germline bias, will be necessary to fine-tune global vaccination efforts and delineate missing genetic risk factors in autoimmune disease ^12–16^.

The human IGH locus is found at the telomeric end of the q arm of chromosome 14 and spans approximately 1Mb. It encodes approximately 54 functional IGHV genes, 28 IGHD and 6 IGHJ genes, in addition to numerous pseudogenes and the IGH constant genes ^4^. The distance encompassed by the IGH locus varies due to the frequent occurrence of structural variations that result in differences of several hundred kb between haplotypes ^17^. Commonly observed structural variations include contiguous deletions that encompass one or more IGHV genes, such as the deletions of IGHV1-8 and IGHV3-9, IGHV5-10-1 and IGHV3-64D, as well as the deletion of six contiguous IGHD genes between IGHD2-2 and IGHD3-9 ^3,18^.

Gene duplications are also common, including a segment including IGHV1-69, IGHV1-69-2 and IGHV2-70, the IGHV3-30 associated IGHV genes, and a separate segment including IGHV3-23. Several structural variants are present at a high frequency in the population, with most individuals having one or more variants on their diploid IGHV locus and many being homozygous for specific variants ^4^. In addition, IGHV allelic diversity and heterozygosity is common. Furthermore, many IGHV genes have common allelic variants that are utilized at widely different frequencies within the naïve repertoire ^19^. Thus, inter-individual variations in IGH genotypes greatly impact the composition of the expressed B cell repertoire ^3,20^.

While the heavy chain (HC) complementarity-determining region 3 (HCDR3), which spans the HC VDJ junction, is a major determinant for epitope recognition for most antibodies, many paratope-epitope interactions involve germline encoded sequences both within and external to the heavy and light chain CDR3s. Since the CDR1 and CDR2 are entirely germline-encoded, inherited variations in IG genes, particularly surrounding the CDRs, have the potential to profoundly influence multiple biological processes. Indeed, antibodies that display virus neutralizing capacity prior to SHM-associated affinity maturation ^21,22^ or following very limited SHM ^23,24^, were described. Furthermore, a growing number of observations have identified antibodies that require a certain IG allele usage ^25–31^, demonstrating the importance of understanding germline-encoded IG variation, not least to guide germline-targeting vaccine strategies ^32^. Delineating the extent to which polymorphisms in IG genes influence responses in a variety of physiological settings is likely to be a complicated process, even limiting this to currently known pathogens. However, the initial step required, that of acquiring a comprehensive understanding of global human IG gene variation, is clear.

Unlike most genomic regions, the IGH locus is refractory to accurate mapping using short read (<300 bp) sequencing approaches. Its location, in the immediate vicinity of the telomere of chromosome 14q, is within a region that has been subject to segmental duplications and deletions during evolutionary times producing additional functional genes as well as the gradual expansion of non-productive but highly similar pseudogenes. These features render short-read sequence-based bioinformatic approaches highly problematic ^33,34^. Longer-read approaches using bait-capture Pac-Bio sequencing have proven more successful as they have sufficient contextual sequence to distinguish the correct genes ^3,17^. However, long-read approaches have several drawbacks, such as the requirement for large amounts of DNA template for accurate analysis, and the number of cases that can be simultaneously sequenced, thereby limiting throughput.

We previously developed an expression-based BCR and TCR genotyping approach, named IgDiscover ^35,36^. While highly accurate and reproducible, expression-based genotyping requires a source of B cells, usually peripheral blood mononuclear cells (PBMC) and it is therefore not easily scalable. We surmised that an approach that can accurately genotype the set of functional IGHV, IGHD, and IGHJ genes in a multiplex manner, while using minimal quantities of DNA template, could be utilized to analyze germline-encoded variation in large cohorts of individuals. Such an approach would facilitate the analysis of previously collected sample sets, such as those representing diverse human populations, and from disease cohorts, allowing association studies of variation within the adaptive immune receptors. To meet these critical requirements, we developed a high-throughput BCR/TCR genotyping approach, ImmuneDiscover. The approach uses nanogram quantities of template DNA as the basis of a targeted multiplex amplification of unrearranged germline sequences within a 96-well format, resulting in an individual library of locus-specific sequences within a single well. The ImmuneDiscover method enables the indexing and simultaneous sequencing of approximately 50 target genes from 1000 individuals per MiSeq run, producing full IGH genotypes from each case, as well as identifying novel variants.

Here, we applied ImmuneDiscover to define IGH variation at the global population level using a set of 2486 samples from the 1000 Genomes Project (1KGP) set, comprising 25 population groups. We identified 561 IGHV allelic variants of functional IGHV genes in this set, thereby enabling accurate frequency estimations of global gene and allele variation within this locus. We found large population biases in the presence of many gene variants and homozygous gene deletions that are critical for the response to numerous pathogens and highly relevant to ongoing clinical vaccination studies ^37^. Upon peer-reviewed publication of this manuscript, the data will be available as an open access resource that allows analysis of sequences and population frequencies, the Karolinska Institutet Adaptive Immune Receptor Gene Variant Atlas (KIARVA), for future studies of adaptive immune receptor gene variation.

## Results

### Development and benchmarking of ImmuneDiscover

An adaptive immune receptor genotyping approach that facilitates the analysis of high numbers of samples requires both accuracy and reproducibility. Such a genotyping technique should, additionally, require a minimal amount of template genomic DNA so that it is broadly applicable to clinically relevant samples. Furthermore, the development of a high throughput genotyping technique requires validation of the output through comparison with an independent, well-established methodology used on the same sample set. To achieve this goal, we developed ImmuneDiscover and performed extensive benchmarking using a validation cohort comprising 90 individuals, herein referred to as the KI cohort, from which both PBMCs and genomic DNA samples were available. Each donor of the KI cohort self-identified as a member of one of four continental population groups, namely African, East Asian, European and South Asian (**Figure 1A**).

**Figure 1.**
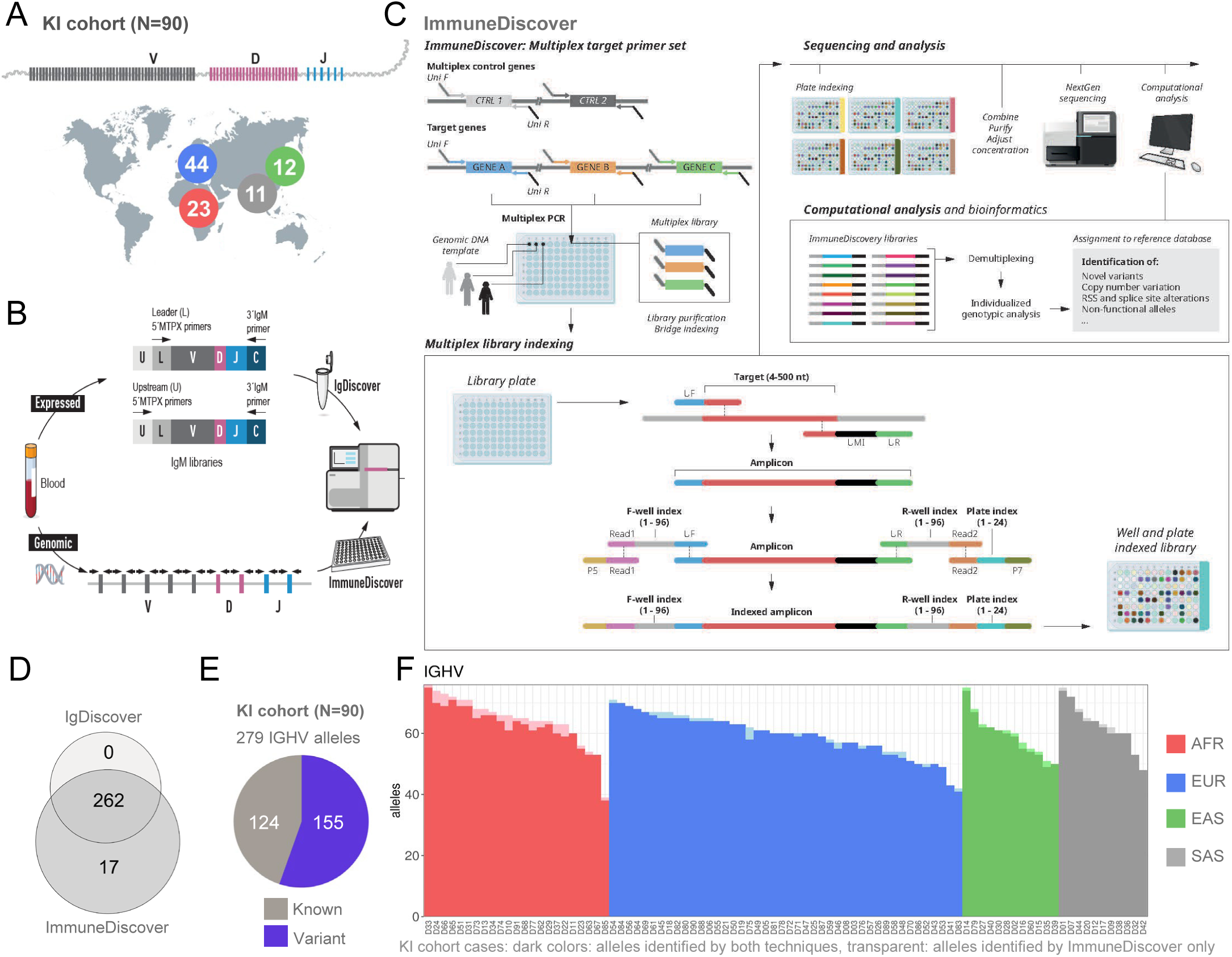
IGH genotype analysis of the KI cohort samples and technique validation. **(A)** Schematic of the genomic organization of the IGH locus consisting of non-rearranged IGHV, IGHD and IGHJ germline genes and of the 90 individuals of African, European, East Asian or South Asian origin who provided a blood samples. **(B)** Overview of the IG genotyping cross-validation process. PBMCs from each case were used for RNA isolation for IGM library preparation. Two independent 5’ multiplex-based libraries were created using primer sets located within the IGHV gene leader region (L) or 5’ UTR region (U), each of which enabled the production of full length VDJ libraries. Following MiSeq sequencing, IgDiscover and corecount analyses were performed to produce individualized IG genotypes for each case. Simultaneously, genomic DNA from each case was used as template for targeted amplification of IGHV, IGHD and IGHJ genes using the ImmuneDiscover technique, followed by multiplex indexing and analysis. **(C)** The ImmuneDiscover analysis method. Left upper panel, each functional gene was targeted by a gene-specific primer pair that contains 5’ Universal forward (Uni F) and Universal Reverse (Uni R) tails. 96-well PCR plates containing genomic DNA from single individuals in separate wells. Primer sets that amplify each target gene set (IGHV, or IGHD plus IGHJ) and control genes, to produce multiplex libraries in each well. Each target amplicon contains Uni F and Uni R tails at the 5’ and 3’ ends of each sequence that are utilized in the indexing procedure. The indexing process uses bridging primers to introduce F and R well-specific indexes as well as a plate index to each separate library to enable combination of libraries from multiple plates for sequencing and subsequent demultiplexing based on the well and plate index content of each amplicon. The ImmuneDiscover software performs the demultiplexing, identifies variant germline alleles and identifies copy-number variation to produce an individual IG genotype. **(D)** Venn diagram showing overlap of IGHV alleles identified using IgDiscover compared to those identified from genomic DNA using ImmuneDiscover. **(E)** Pie chart of the IGHV alleles identified in the 90 cases, showing the number of alleles present in the AIRR-C reference database (Known) and the number of alleles that were not in the starting database (Variant). **(F)** Stacked bar chart of the number of IGHV alleles present in each of the 90 donor cases with the alleles found solely by the ImmuneDiscover shown in a lighter shade for each case.

We utilized the germline allele inference program IgDiscover to produce individual IG genotypes from expressed IgM repertoires from each of the 90 KI cohort cases for cross-comparison with the IG genotypes generated by genomic DNA typing using ImmuneDiscover (**Figure 1B**). For the IgDiscover analysis, two IgM libraries, using independent 5’ multiplex PCR primer sets located in the 5’ UTR region or the leader region, respectively, of all functional IGHV genes, were produced for each case, as described previously ^36,38^, thus entailing the production, sequencing and analysis of 180 IGM libraries in total. The use of two independent primer sets enabled improved coverage of genes that have rare sequence variation affecting 5’ primer binding. Each IgM library was sequenced on the Illumina MiSeq platform and analyzed using the germline inference tool IgDiscover, and the genotyping tool corecount ^39^ to produce an IGHV, IGHD and IGHJ genotype for each case. We utilized the AIRR-C reference set ^10^ as a starting database for the analysis to both confirm previously well-validated alleles and enable the identification of additional variants.

We next performed genomic DNA-based IGH typing of the KI cohort cases using the ImmuneDiscover technique, which relies on targeted multiplex library amplification of the members of a multigene locus within the wells of a 96-well plate that contains template DNA from a single individual. This is followed by a highly specific procedure that enables the indexing of each library with two well-specific indexes and a plate-specific index. The indexing procedure utilizes a bridge PCR approach to simultaneously add both the well and plate indexes (**Figure 1C**). This combination of indexes enables the libraries from multiple plates, each containing up to 94 test libraries along with negative and positive controls, to be combined and sequenced together, with the individual libraries subsequently demultiplexed by the ImmuneDiscover software. The program utilizes a BLAST-based process to identify new allelic variants. The libraries are analyzed individually, enabling the identification of gene variants, recombination signal sequence (RSS) and splice site variants, and copy number variants, with the resultant output being a locus genotype for each case (**Figure S1**). The ImmuneDiscover process utilized 4 ng of DNA template to amplify the set of IGHV and the set of IGHJ and IGHD allelic variants from each of the 90 cases using the appropriate multiplex primer sets. Primer sets specific to either the IGHV or the IGHD plus IGHJ genes were utilized to simultaneously amplify products of approximately 450 bp in a 96 well plate, thereby producing libraries of similar length targets that can be sequenced using the Illumina MiSeq instrument and subsequently processed by the ImmuneDiscover software. To test the reproducibility of the ImmuneDiscover analysis, we generated 16 separate libraries each from 3 cases from the KI cohort. The results revealed identical individual genotypic outputs from almost all 16 libraries for each case, with just 1 of 15 replicate libraries from cases D19 and D25 showing zero false positive alleles and just a single false negative allele (**Figure S2A**).

A total of 262 allelic variants of functional IGHV genes were identified using IgDiscover, while 279 IGHV alleles were identified through genomic DNA analysis by ImmuneDiscover. Critically, all 262 alleles identified from the expressed IgM libraries by IgDiscover were also identified by ImmuneDiscover (**Figure 1D**). Further analysis of the 17 alleles identified only with ImmuneDiscover revealed that they were non-expressed alleles (**Figure S2B**). These included known non-functional allelic variants of functional IGHV genes, including IGHV3-20*02 and IGHV6-1*03, in addition to variants of IGHV3-49 and IGHV3-53 that contained stop codons due to indels. Of the 279 alleles, 155 alleles were not present in the starting reference AIRR-C database (**Figure 1E**). Importantly, the exact same alleles were identified by both IgDiscover and ImmuneDiscover in each person (**Figure 1F**, dark colors), except for the non-expressed alleles that were only identified with ImmuneDiscover (**Figure 1F**, transparent colors). The total number of IGHV alleles in each of the 90 cases varied between 40 and 78, depending on heterozygosity and/or the presence of homozygous gene deletions (**Figure 1F**). In addition to the IGHV analysis, 10 IGHJ alleles and 39 IGHD alleles were identified through IgDiscover analysis of the KI cohort.

### IGHV genotypic analysis of the 1KGP cohort

To identify additional germline variation and delineate the allelic frequencies within different global populations, we next extended the ImmuneDiscover analysis to a larger, well-defined population cohort, the 1KGP. A total of 2486 cases from 25 separate populations were analyzed. These included 7 African origin populations of which two (ASW and ACB) were collected in the Americas, 5 South Asian populations, 5 East Asian populations, 4 European populations, and 4 American origin populations (**Figure 2A**). Each of the 1KGP case templates consisted of DNA extracted from immortalized lymphoblastoid cell lines created from EBV transformation of PBMCs. This process poses an issue as PBMC-derived B cells may contain VDJ-associated deletions in cells that have rearranged both IGH haplotypes. As functional VDJ recombination occurs through sequential DJ and then VDJ recombination events, an individual template may contain a mixture of DNA from complete IGH, DJ, and VDJ recombinant haplotypes (**Figure S2C**). ImmuneDiscover genotypic analysis with the combined D and J primer set revealed that most samples contained multiple unique DJ recombinant clones, as shown by the presence of library specific amplicons that utilized D gene 5’ primers and J gene 3’ targets. This indicates the presence of multiple DJ recombinants rather than VDJ recombinants, with concomitant presence of the full IGHV haplotype within each of these DJ clones. The frequency of multiple DJ identification thereby enabled the identification of accurate full IGHV genotypes from most cases. We used the number of IGHV alleles as measure of completeness for each IGHV genotype (**Figure S2D** and **E**). Based on a comparison with the number of IGHV alleles per individual found in the KI donor cohort, we used 35 IGHV alleles as a cutoff and found that 97.9% (2435) 1KGP cases were above this number and thus were suitable for IGH genotypic analysis (**Figure S2F**).

**Figure 2.**
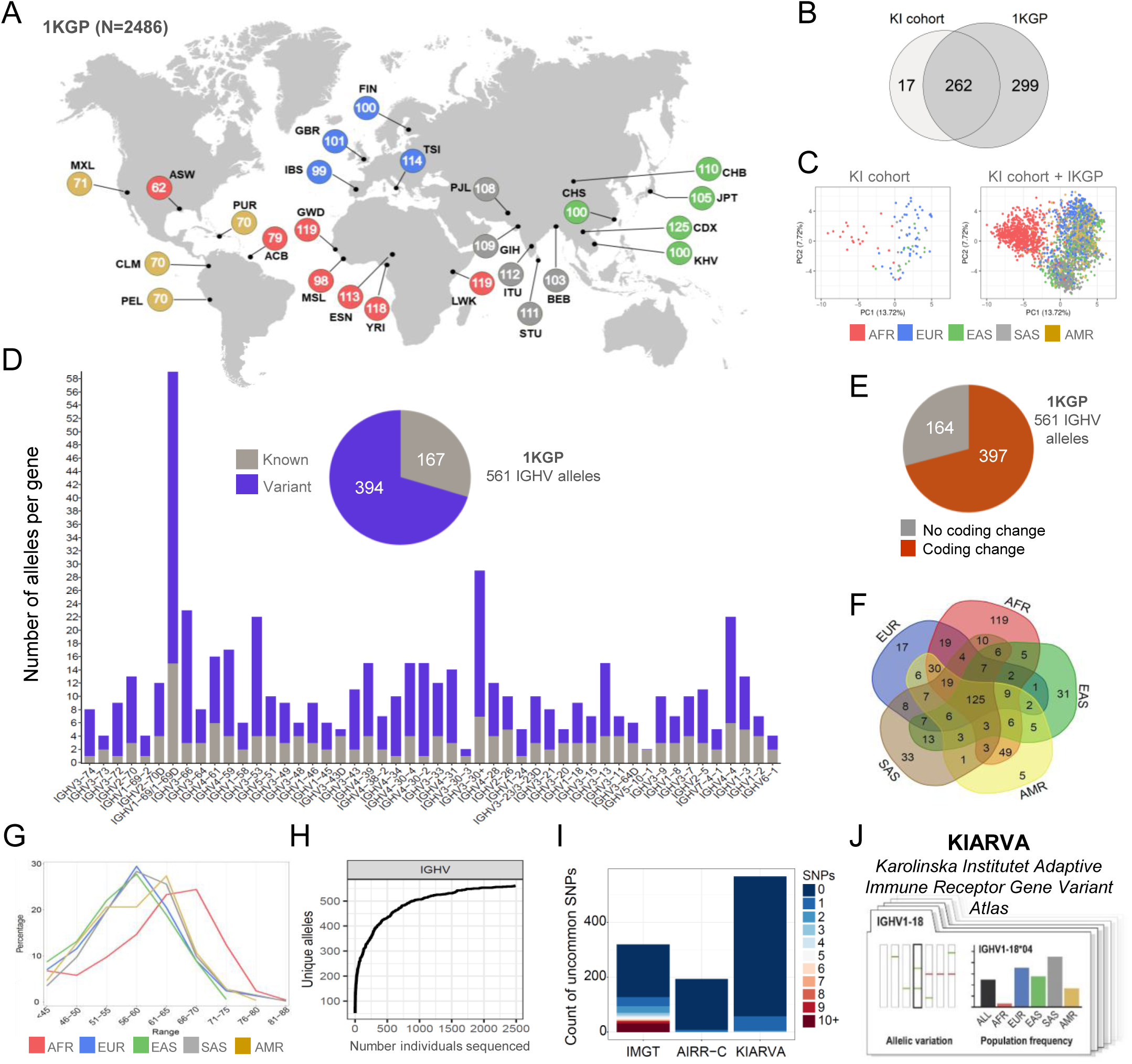
IGH genotype analysis of the 1KGP samples and the KIARVA resource. **(A)** Populations from the 1KGP sample set were IGH genotyped using ImmuneDiscover. The number of cases in each of the 25 population groups is shown within the colored circles, with the total number being 2486. The continental origin-based superpopulation groups, African, European, East Asian, South Asian and American, are indicated in the key. **(B)** Overlap of unique IGHV alleles identified in the KI cohort and the 1KGP set using ImmuneDiscover analysis. **(C)** PCA analysis of individuals from the KI cohort set (left panel) and the combined KI cohort and 1KGP cases (right panel) based on their IGHV genotypes. Each case is illustrated by a point, colored based on the superpopulation group. **(D)** Number of alleles per IGHV gene found in the 1KGP dataset. Known alleles present in the AIRR starting database are shown in grey, and previously unknown (variant) IGHV alleles are shown in purple. The total numbers of known and variant IGHV alleles are shown in the inset pie chart. **(E)** Pie chart illustrating the number of IGHV alleles with coding changes of the total 1KGP set of 561 alleles. **(F)** Overlap of IGHV alleles present in individuals from each of the five 1KGP superpopulations. **(G)** The range of allele counts found in each genotype for each superpopulation. **(H)** Saturation curve analysis of the number of unique IGHV alleles identified in the 1KGP for each genotype analyzed. **(I)** Stacked bar chart illustrating the results of IgSNPer analysis of all IGHV alleles present IMGT), AIRR-C and KIARVA reference databases with high IgSNPer score indicating alleles that were not substantiated by available SNP databases. The IgSNPer score key is shown in the key on the right of the panel. **(J)** Generation of the KIARVA resource comprising 1KGP IG alleles and population frequencies.

ImmuneDiscover analysis of the 1KGP cohort revealed a total of 561 allelic variants of functional IGHV genes, of which 262 were also present in the KI donor cohort. 17 variants found in the KI donors were not present in the 1KGP genotypes (**Figure 2B**), each of which were present at low numbers in the 90 donors. We next compared the IGHV alleles identified in the 1KGP superpopulation groups, defined as the combined population groups of the same recent continental origin, with the 90 KI donor group using principal component analysis (PCA) analysis. We found that the KI cohort clustered similarly to the 1KGP set, with African cases being the most distinct (**Figure 2C)**. Additional variants were identified for each functional IGHV gene, compared to the known IGHV alleles described in the starting database. In total, 394 of the 561 IGHV alleles were identified in the 1KGP set (70%) were missing from the

AIRR-C reference database (**Figure 2D**). Of the 561 IGHV alleles present in the 1KGP set, 397 variants contained amino acid coding variations and 16 are non-functional due to internal frameshift mutations or stop codons (**Figure 2E**).

Individual genotyping of each of the 1KGP donors enabled the identification of the sets of allelic variants present for each population and superpopulation (**Figure 2F**). As expected, there was a large overlap, with 125 alleles found in all 5 superpopulations. The African superpopulation contained the highest number of IGHV alleles (n=119) that were only found in that group, followed by the 33 alleles in the South Asian group, 31 alleles in the East Asian group, 17 alleles in the European group and 5 alleles in the American group (n=5) (**Figure 2F**). Analysis of functional IGHV allele counts per individual demonstrated a range of allele numbers in the populations, with the highest numbers of alleles found in the African superpopulation due to a higher degree of IGHV heterozygosity and gene duplications in this population **(Figure 2G).**

A key objective of the current study was to identify a substantial fraction of the most frequent global IGHV variants, enabling the construction of a validated reference set for future IG analysis studies. We therefore performed a saturation analysis of the alleles, based on the number of unique alleles present in each additional case. The result indicated that we approached saturation for the populations in the 1KGP dataset (**Figure 2H**). As indicated by the additional variants present solely within the KI cohort, additional variants will be found as more genotypes are analyzed: however, these are likely to be rare, at least within the population groups analyzed in the current study.

### Validation of datasets and utilization of IgSNPer to include large population SNP datasets

IG alleles identified within the KI cohort were cross validated by comparing the results of the two independent IgDiscover libraries, inferred-haplotype analysis and ImmuneDiscover analysis for each case. To ensure the accuracy of the alleles identified by ImmuneDiscover in the 1KGP samples, we utilized a cross-validation approach that involved the alleles to be present in more than one case.

While the IG genotypic analysis of several thousand individuals provides a prime mechanism to identify germline frequencies within the populations included, analysis of even larger human datasets provides an additional layer of cross-validation of the variants identified. To achieve this, we utilized the availability of datasets comprising up to 1.25 million individuals genotyped for genome-wide single nucleotide polymorphism (SNP) variants, a selection of which we could localize to gene-specific IG alleles. This enabled the creation of a bioinformatic analysis tool named IgSNPer. This software collates a series of independent population datasets (**Figure S3**) containing the results of SNP variant analysis from six databases involving approximately 1.25 million individuals worldwide. Since SNP variants are commonly mapped to two genomic assemblies, GRCh37/hg19 and GRCh38/hg38, we utilized these assemblies to cross-map the genomic coordinates of the genotyped alleles, thereby identifying the frequency of specific SNP variants located within allele-specific variants. The tool enables the testing of each allele specific variant SNP that can be mapped to an assembly coordinate against the IgSNPer variant database, resulting in an IgSNPer score of 1 for each SNP that falls below the cutoff frequency for the program. An IgSNPer score of zero indicates that each variant nucleotide present within that sequence is also present in multiple individuals in the IgSNPer SNP variant database. A low score thereby indicates high confidence, while a higher score indicates lower confidence of the variant being represented within the 1.25 million case collated SNP dataset.

The program can be utilized to examine collections of alleles, such as those present within reference databases, identifying variants that may be erroneous due to truncations, sequencing errors or accidental inclusion of sequences containing somatic hypermutation, each of which is expected to result in a high IgSNPer score. In the case of the 1KGP allele set produced in this study, the vast majority of alleles that could be mapped to an assembly produced an IgSNPer score of zero, with a small percentage showing scores of 1 or 2. In contrast to this, analysis of the IMGT reference database indicated that many variants present produced high IgSNPer scores, mainly, although not only, due to substantial truncation of allele length. In contrast, both the well-validated, albeit size-limited, AIRR-C IG database and the KIARVA database generated here showed a low number of alleles with IgSNPer scores above zero (**Figure 2I)**. The full-length high IgSNPer score alleles present in IMGT were absent from both the KI cohort and the 1KGP set. Sequences with clear sequence errors or ambiguities, for example IGHV1-2*03, suggest that erroneous variants are present at a non-negligible level in the commonly used IMGT reference set. Crucially, IgSNPer functioned as an independent validation for newly identified germline alleles during this study, enabling the expansion of case number analysis from several thousand to more than 1.25 million individuals.

To make the IG gene variation identified in the 1KGP set available to the research community we created an open access resource: KIARVA. This resource comprises downloadable FASTA files and evidenced IGHV population frequencies across the 25 subpopulations in the 1KGP set, as well as sequence alignments at the nucleotide and amino acid levels (**Figure 2J**). Thus, the KIARVA resource provides both an opportunity to identify critical coding differences that may affect paratope-epitope interactions and enables an understanding of baseline frequencies of these variants in multiple global population groups.

### Distinct patterns of IGHV structural variation at the population level

Since structural variation is a major source of inter-individual IGH variation, we utilized ImmuneDiscover to first produce individualized genotypes for the 90 KI cohort cases to enable analysis at the gene level. The allele number per gene, or absence thereof, allowed us to identify the level of homozygosity, heterozygosity, duplication, and deletion for each gene (**Figure 3A**). Structural variation was common at both the individual and population level in the 90 donors of the KI cohort, with common IGHV deletion regions, such as of IGHV2-70D and IGHV1-69-2, IGHV3-43D, IGHV4-38-2, IGHV4-30-4 and IGHV4-30-2, IGHV3-33 and IGHV4-31, IGHV3-64D and IGHV5-10-1 and IGHV 3-9 and IGHV1-8, in multiple populations.

**Figure 3.**
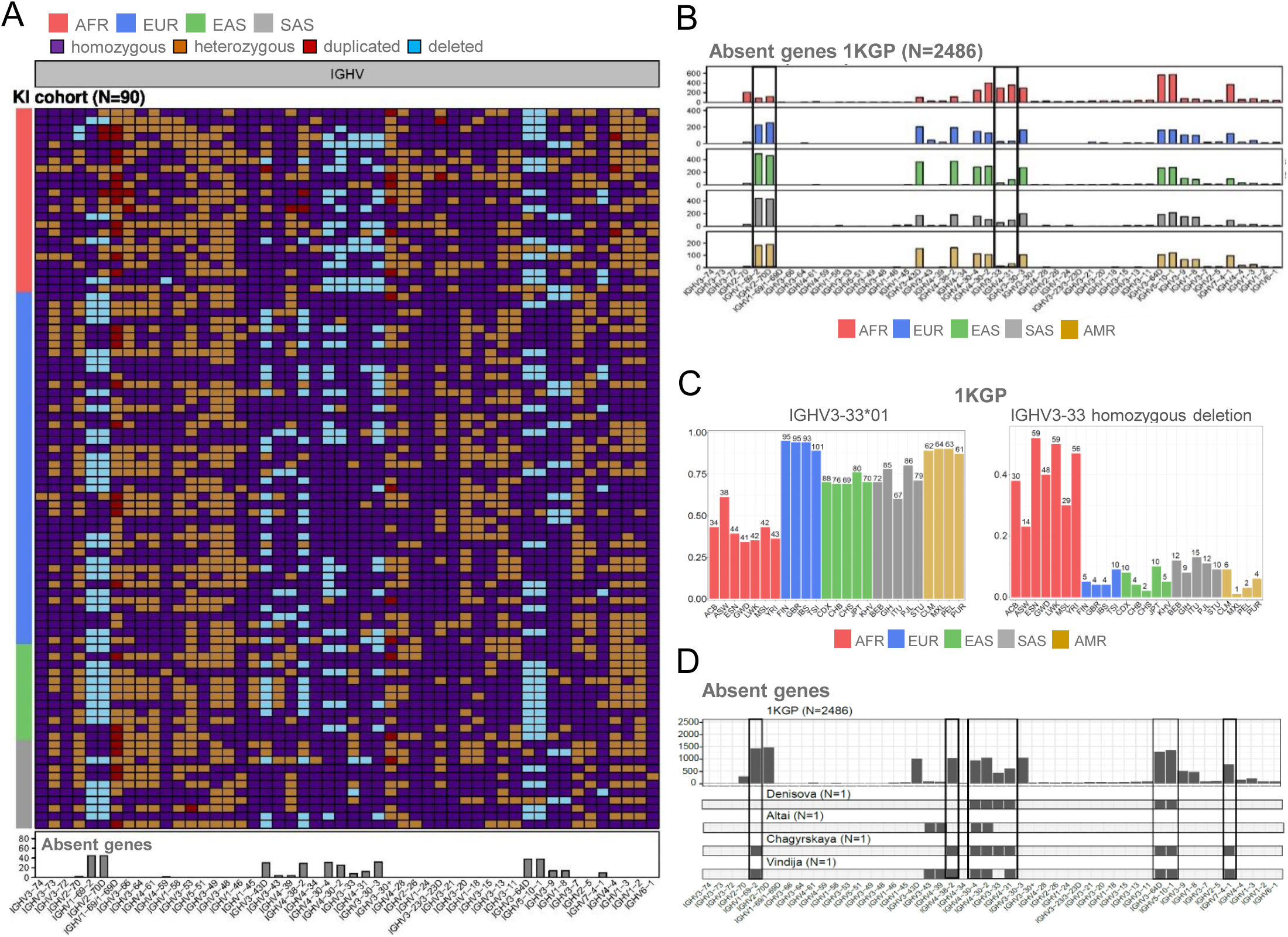
Population-based analysis of IGHV gene zygosity and gene deletions. **(A)** IGHV gene status in the 90 KI cohort cases (y-axis) showing homozygous presence (dark blue), heterozygous (orange), duplications (red) and homozygous deletions (light blue) with the gene names listed in chromosomal order (x-axis). Genes with allelic variants that are identical at the nucleotide level were combined as IGHV1-69/IGHV1-69D, IGHV-3-23/IGHV3-23D and IGHV3-30/IGHV3-30-5 (labeled IGHV3-30+), respectively. Summary of homozygous IGHV deletions (Absent genes) in the KI 90 cohort is shown below the tile plot. **(B)** Number of individuals with homozygous IGHV deletions in each of the 5 superpopulation groups in the 1KGP set. The IGHV1-69-2-IGHV2-70D and IGHV3-33– IGHV4-31 structural variants that show the greatest variance in different populations are indicated by the boxed areas. **(C)** Example of KIARVA output plots showing IGHV3-33 variation within the 1KGP subpopulations where the upper panel shows the frequency of the IGHV3-33*01 allele and the lower panel shows the frequency of homozygous IGHV3-33 deletions. **(D)** IGHV gene deletion analysis in high coverage assemblies of archaic humans compared to the 1KGP set. The upper panel shows the combined results for the 2486 individuals from the 1KGP set and the lower panel shows IGHV gene deletions in the Altai, Chagyrskaya and Vindija Neanderthal assemblies and the single high coverage Denisovan assembly. Commonly absent genes within the archaic individuals are shown in the boxes.

We next analyzed the 1KGP set, which showed that these deletions are also common in the larger dataset. Critically, this analysis demonstrated that the frequency of specific gene deletions varied by superpopulation. For example, the African 1KGP cases showed frequent absence of the contiguous set of IGHV4-30-4, IGHV4-30-2, IGHV4-31 and IGHV3-33, while having the lowest level of deletion of IGHV1-69-2 and IGHV2-70D. In contrast, IGHV1-69D and IGHV2-70D are absent at the highest frequency in East Asian cases, while absence of IGHV1-8 and IGHV3-9 is highest in European cases (**Figure 3B**). Analysis of the number of IGHV alleles per person in the 1KGP sample set is shown in **Figure S4**. Homozygous IGHV3-33 gene deletions may be especially relevant in the context of vaccine efforts aimed at re-eliciting a class of antibodies that target the *Plasmodium falciparum* circumsporozoite protein (PfCSP), which utilize IGHV3-33 ^40,41^. The ImmuneDiscover results shown here indicate that this gene is frequently absent in the African superpopulation group, with over 40% of some African populations having homozygous loss of this gene (**Figure 3C**).

Population differences at the IGHV allele level were also apparent with one example being IGHV6-1, a gene used in multi-donor broadly neutralizing influenza antibodies ^42–45^ (**Figure S5A**). We found that heterozygosity of IGHV6-1 was more frequent in individuals of African ancestry where both the IGHV6-1*01 and IGHV6-1*03 alleles were common, in contrast to other population groups where IGHV6-1*01 is the predominant allele. IGHV6-1*03 encodes a premature stop codon, rendering it non-functional (**Figure S5B**). In the 1KGP samples, we found that approximately 18% of the African population was homozygous for IGHV6-1*03, indicating absence of this gene in the expressed B cell repertoire of these individuals (**Figure S5C**).

We previously performed genotypic analysis of high coverage archaic genomic assemblies, revealing introgression of Neanderthal TCR gene variants into modern human population ^35^. The IGH locus, in contrast, contains many more pseudogenes rendering accurate allelic genotyping far more difficult since high coverage archaic sequencing necessitates short read sequencing. These short sequences can be highly problematic to assemble accurately due to the high similarity between functional genes and non-functional pseudogenes. We found, however, that there is sufficient sequence identity at the gene, rather than allele level, to identify the presence of archaic sequences corresponding to most human functional IG genes. This enabled the production of IGHV gene presence results for the three Neanderthal assemblies from the Altai, Chagyrskaya and Vindija individuals and from the single Denisova high coverage assembly ^46–49^, revealing that several of the structural deletions common in modern populations were homozygous in the archaic individuals (**Figure 3D**). The more recent surviving archaic samples, namely the Chagryskaya and Vindija Neanderthals, contained the lowest number of IGHV genes (**Figure 3D** and **Figure S2G**), with homozygous deletion of 9 and 10 IGHV genes respectively.

### IGHJ and IGHD structural and allelic variation between population groups

Genetic variation of IGHD and IGHJ genes is also identifiable between individuals and populations. These genes are frequently subjected to DJ recombination in lymphoblastoid cell line material that results in partial loss within the IGHD and IGHJ regions of the locus. Thus, care must be taken to ensure accuracy of individual genotypes. Most IGHD and IGHJ variants described in this study were identified using both IgDiscover and ImmuneDiscover library analysis from the 90 donors of the KI cohort. These additional variants were also confirmed through ImmuneDiscover analysis to be present within the 1KGP set. Overall, IGHD variation is more extensive than IGHJ variation, at both the allele and gene deletion level (**Figure 4A**).

**Figure 4.**
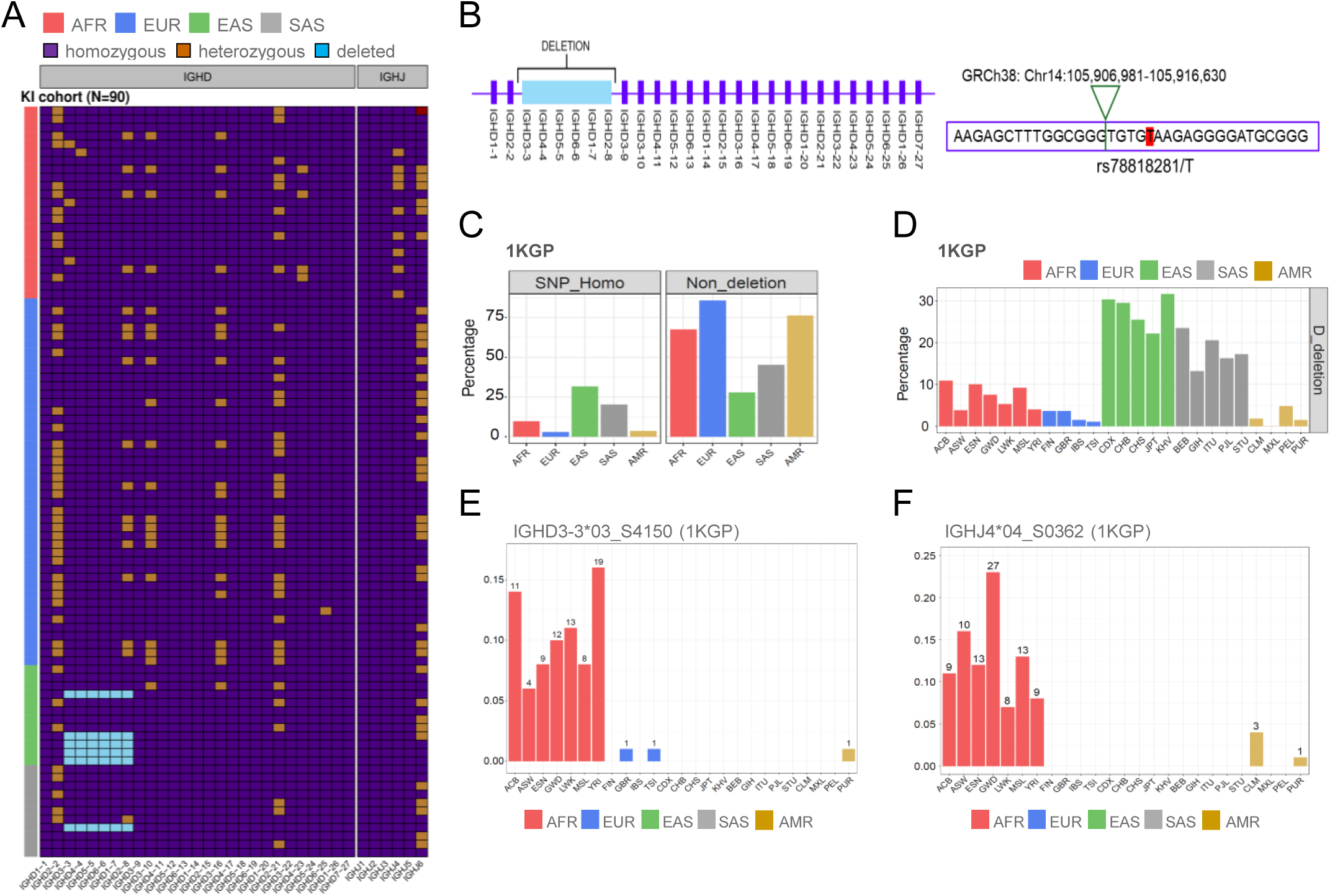
Structural variation in the IGHD locus in the KI cohort and 1KGP sample sets. **(A)** Tile plot illustration of IGHD and IGHJ gene variation in the 90 KI cohort cases showing homozygous presence (dark blue), heterozygous (orange) and homozygous deletions (light blue). **(B)** Schematic of the deletion that removes six IGHD genes between IGHD2-2 and IGHD3-9 is shown to the right with the deletion breakpoint-associated SNP variant rs78818281/T shown to the left. **(C)** Frequency of 1KGP superpopulation cases that show homozygosity for the rs78818281/T deletion variant or homozygosity for the non-deletion variant, rs78818281/G. (D) Frequency within the 1KGP populations showing genotypic presence of the retained IGHD2-2 and IGHD3-9 genes in conjunction with absence of all six genes, IGHD3-3, IGHD4-4, IGHD5-5, IGHD6-6, IGHD1-7 and IGHD2-8 within the deletion haplotype. **(E)** Presence of the IGHD3-3*01_S4150 variant within the 1KGP population groups. **(F)** Presence of IGHJ4*04_S0362 variant within the 1KGP populations.

ImmuneDiscover analysis revealed the presence of a known IGHD deletion haplotype in multiple populations. This deletion involves the loss of a contiguous set of 6 IGHD genes between IGHD2-2 and IGHD3-9, with frequent homozygous loss found in individuals of East and South Asian origin in the KI cohort (**Figure 4A**). PCR amplification across the deletion using primers located close to IGHD3-9 and IGHD2-2, in a homozygous deletion case enabled the identification of a SNP rs78818281/T that was present in all deletion cases analyzed (**Figure 4B**). This SNP variant is located within 200 bp of IGHD2-2, which was present within the ImmuneDiscover IGHD2-2 targeted amplicon. IGHD2-2 analysis using ImmuneDiscover therefore enabled the typing of this SNP variant, allowing for analysis of IGHD gene content by cross-comparison with the rs78818281/T SNP variant. Using this approach, we found that hemizygosity and homozygosity for the multigene IGHD deletion was present in all superpopulations but was particularly high in East Asians, with South Asians a close second (**Figure 4C** and **D**). We identified several previously undescribed IGHD variants, mostly from the KI cohort cases (**Figure S5D**). This included IGHD3-3*03_S4150, which was found in all African 1KGP subpopulation groups (**Figure 4E**). Since IGHD genes can be utilized in multiple reading frames, the presence of additional IGHD alleles presents opportunities for a greatly increased range of functionality within the recombined HCDR3 repertoire (**Figure S5E**). Indeed, several well-characterized broadly neutralizing antibodies against influenza HA stem utilize IGHD3-3, often with IGHV6-1. For these antibodies, our results from the 1KGP samples suggest that population restrictions may arise from either a lack of functional IGHV6-1, the IGHD3-3 deletion described above, or both ^50^. These results are also relevant for HIV and SARS-CoV-2 that were also shown to utilize specific IGHD genes and frames ^44,51–53^. Allelic variation in IGHJ genes other than IGHJ6*02/*03 is rare, except for IGHJ4*04_S0362, a variant of IGHJ4 which was found in over 10% of African cases (**Figure 4F** and **Figure S5F**).

### IGHV variations have consequences for antibody functionality

Analysis of IG gene variation at the nucleotide level is important in defining population structure; however, it can potentially obscure functional similarities. Collapsing IGHV nucleotide variants to their translated amino acid sequences enables functional comparisons of population-biased variants that encode identical translated sequences. Additionally, it enables the delineation of germline variants containing HCDR1 and HCDR2 coding differences that may directly affect paratope functionality. We therefore set up the KIARVA resource to allow comparison between allelic variants of each gene both at the nucleotide level and as collapsed amino acid sequences, in conjunction with the population frequency of each. A total of 29/51 functional IGHV genes include germline HCDR1 variants, 29/51 HCDR2 variants and 6 included HCDR3 region coding variants. In addition, 16/51 IGHV genes had coding variation with the IGHV framework 3 region, between amino acids 70-78, a segment equivalent to the TCR hypervariable 4 (HV4) region, indicating the possibility that variation in this region of some IGHV genes may have functional consequences. Finally, two IGHV genes, IGHV4-34 and IGHV5-10-1, contained germline sequences within their HCDR2 that included motifs for N-glycosylation that could potentially affect their function ^54^.

Analysis of the ImmuneDiscover IGHV libraries additionally enabled identification of common leader and RSS heptamer variants that flanked the IGHV coding sequence. This confirmed the presence of a non-functional IGHV4-34*01 RSS heptamer variant, CTCAGTG, previously identified during an IGHV genotype study of a cohort from Sweden ^55^. The 1KGP results reveal the highest frequency within the Finnish cases, indicative of a higher frequency in Northern European populations. In general human non-functional RSS heptamer and V exon splice site variation is infrequent within the IGHV locus, in contrast to the more frequent variation of this kind previously shown to affect human TCR expression ^35^.

Differences in inter-allelic functionality have implications for ongoing HIV vaccination efforts. Several recent studies have shown promising results regarding the targeted expansion of VRC01-class B cell precursors during vaccination trials ^56–58^. This strategy was designed to elicit IGHV1-2-using antibodies mimicking the VRC01 class of broadly neutralizing antibodies found in some people living with HIV. Critically, the IGHV genotype is a crucial component of the process since only IGHV1-2 alleles bearing the HCDR2-located amino acid W50 are permissive for the VRC01 binding mode ^59^. In this study, we identified seven IGHV1-2 allelic variants within the 1KGP dataset (**Figure 5A),** of which six had unique coding sequences (**Figure 5B**). Analysis of the frequency of these alleles in the 1KGP population revealed that the non-permissive IGHV1-2*05 allele is frequent within the East and South Asian superpopulations (**Figure 5C**). In particular, the frequency of individuals that lack permissive IGHV1-2 alleles is above 20% in both populations (**Figure 5D**). We and others have shown in 4 separate clinical vaccine trials that eOD-GT8 60mer and BG505 SOSIP.v4.1-GT1.1 immunogens can bind B cells that utilize either IGHV1-2*02 or *04, thereby enabling expansion of VRC01 precursors, while B cells expressing IGHV1-2*05 or *06, which lack W50, were not engaged in the response ^32,56–58^.

**Figure 5.**
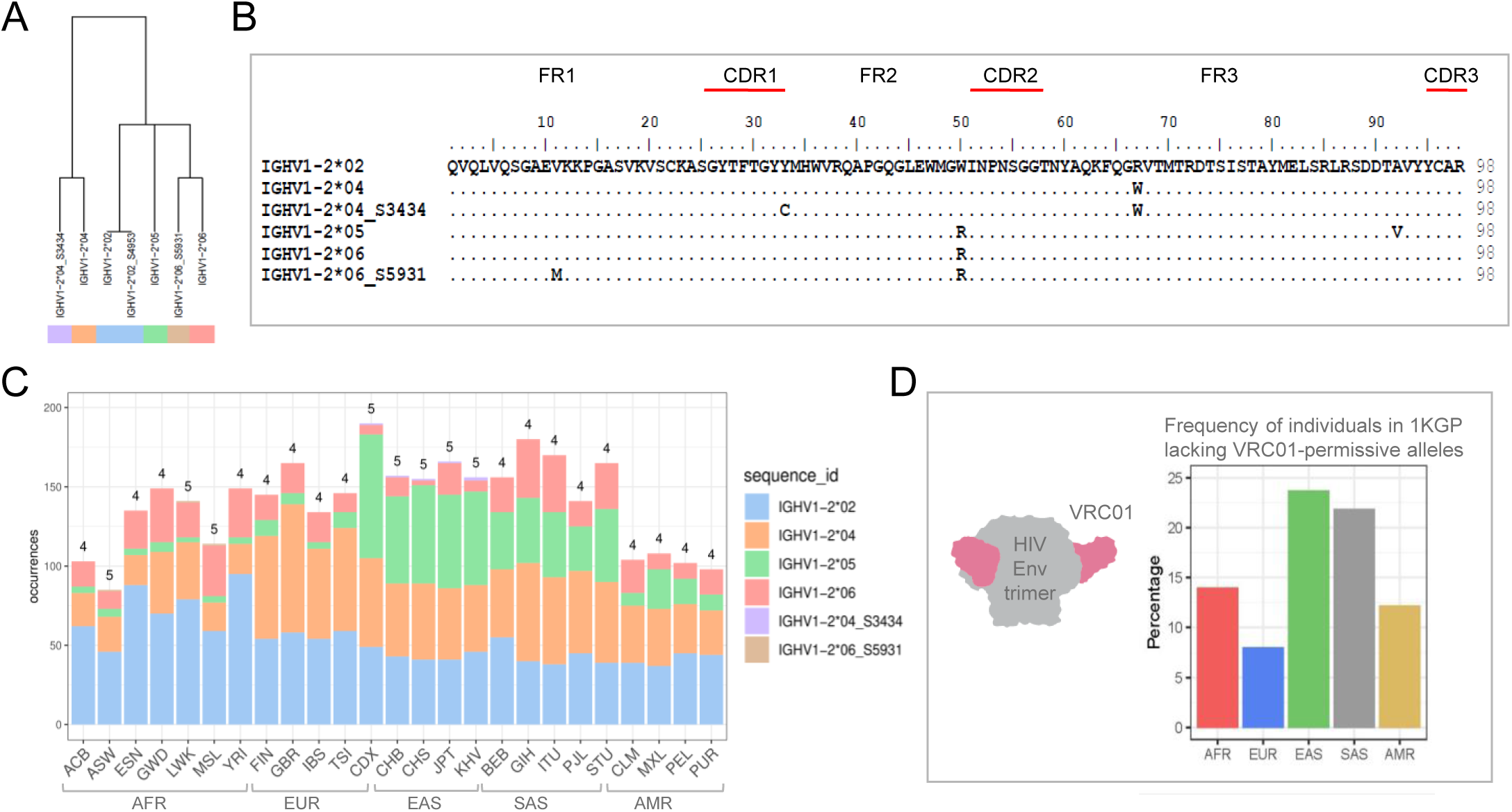
IGHV1-2 variation and allele frequency in the 1KGP populations. **(A)** Dendrogram of the IGHV1-2 alleles identified in the 1KGP set. **(B)** Amino acid alignment of the IGHV1-2 alleles with different coding sequences. IGHV1-2*02, IGHV1-2*04, IGHV1-2*04_S3434 are permissive for the VRC01 antibody class while IGHV1-2*05, IGHV1-2*06 and IGHV1-2*06_S5931 are non-permissive due to the presence of an R instead of a W at position 50. **(C)** Stacked bar chart showing the distribution of the IGHV1-2 alleles with different coding sequences in the 25 1KGP subpopulations according to the color-coded key to the right. The numbers at the top of each bar indicate how many IGHV1-69 alleles with unique coding sequences were found in that population. **(D)** Schematic of the VRC01 class of broadly neutralizing antibodies binding to the HIV trimer and bar chart showing the frequency of individuals in each 1KGP superpopulation who lack VRC01 permissive IGHV1-2 alleles.

It is becoming clearer that the presence of specific germline IG alleles can provide a differential benefit in the immunological response to specific pathogens. The ability of unmutated antibodies to exert neutralizing capacity prior to SHM, for example, provides a distinct temporal advantage. Germline-encoded variation, particularly through amino acids located within the CDRs, is critical in this regard. Analysis of the KI cohort and 1KGP sample sets revealed a remarkably high diversity of IGHV1-69 alleles, with as many as 59 nucleotide variants present in the 1KGP donors (each variant was found in at least 4 independent cases). This resulted in 30 unique amino acid-translated sequences (**Figure 6A** and **S6A**). The 10 most common IGHV1-69 alleles with unique coding sequences were distinguished through amino acid differences in HCDR1, HCDR2 or FR3 (**Figure 6B**). Mapping these 10 IGHV1-69 variants to each of the 25 1KGP populations revealed several clear differences in subpopulation frequencies (**Figure 6C**). As mentioned above, some IGH haplotypes encompassing IGHV1-69 and IGHV2-70 comprise a duplication segment that includes three additional functional genes, IGHV1-69D, IGHV1-69-2 and IGHV2-70D. ImmuneDiscover genotyping of the 1KGP cases revealed population differences in the frequency of this duplication through the presence or absence of IGHV2-70D and IGHV1-69-2. The IGHV1-69 and surrounding genes allelic variation and copy number is highest in the African populations in the 1 KGP set. In contrast, both East Asian and South Asian populations show the lowest average numbers of IGHV1-69, IGHV2-70D and IGHV1-69-2 alleles (**Figure 6D**).

**Figure 6.**
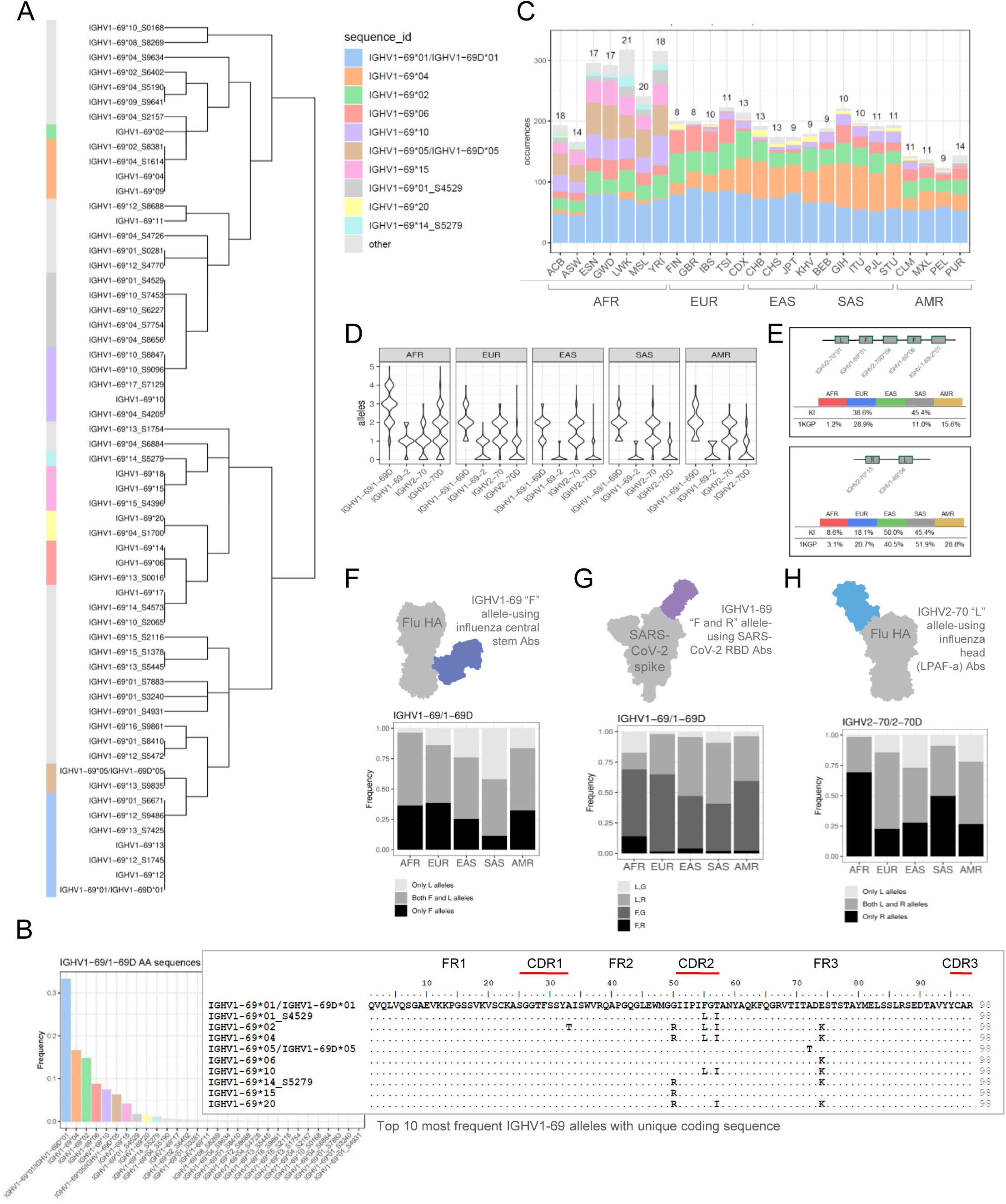
IGHV1-69 variation and allele frequency in the 1KGP populations. **(A)** Dendrogram of the IGHV1-69 alleles found in the 1KGP set. Alleles with nucleotide variation translating to the same amino acid sequence are indicated by the vertical bar and the same color in the left-hand column. The naming key for the ten most frequent IGHV1-69 alleles with unique coding sequences is shown to the right. **(B)** Frequency of IGHV1-69 alleles with unique coding sequences and an alignment of the top 10 most frequent alleles. The lower frequency translated alleles are indicated in grey. **(C)** Stacked bar chart showing the distribution of the ten most frequent IGHV1-69 alleles with unique coding sequences in each of the 25 1KGP subpopulations according to the color-coded key shown in panel A. The numbers at the top of each bar indicate how many IGHV1-69 alleles with unique coding sequences were found in that population. **(D)** Violin plot showing the allele number distribution for IGHV1-69/ IGHV1-69D, IGHV2-70. IGHV1-69-2 and IGHV2-70D in each of the 1KGP superpopulations. **(E)** Schematic of frequent IGHV1-69 region haplotypes. The upper panel shows an IGHV1-69*01/IGHV2-70*01/IGHV1-69*06/IGHV2-70D*04/IGHV1-69-2*01 duplication haplotype and the numbers of cases in the KI cohort and 1KGP set containing this. The lower panel shows a non-duplicated IGHV1-69*04/IGHV2-70*15 haplotype, with the numbers of cases in the KI cohort and 1KGP set containing the alleles from this haplotype. **(F)** Model of IGHV1-69-using anti-influenza HA stem-binding antibody and stacked bar chart of population frequencies of cases containing IGHV1-69 genotypes with only L, only F, or both L and F alleles. **(G)** Model of IGHV1-69-using SARS-CoV-2 RBD-binding antibody and stacked bar chart of population frequencies of cases containing LG, LR, FG or FR alleles. **(H)** Model of IGHV2-70-using LPAF-class anti-influenza HA head-binding antibody and stacked bar chart of population frequencies of cases containing IGHV2-70 or IGHV2-70D genotypes with only L, only R, or both L and R alleles.

Inferred haplotype analysis of the IGHV locus in the 90 donors of the KI cohort revealed the presence of two common IGHV1-69-region haplotypes. One of these, a duplicated haplotype, consisting of two F55-containing IGHV1-69 alleles, IGHV1-69*01 and IGHV1-69*06, and an L52 encoding IGHV2-70*01 allele was common in Europeans, with the allele combination present in approximately 28.9% of 1KGP cases. The other common haplotype we identified here consisted of a non-duplicated IGHV1-69 locus with two alleles, IGHV2-70*15 and IGHV1-69*04. This haplotype was found in 40.5% and 51.9% of the 1KGP East Asian and South Asian cases respectively (**Figure 6E**). The IG haplotype analysis suggests that some individuals contain combinations of IGHV alleles that may provide functionality differences during specific pathogenic exposures. Some of this variation was noted previously as being critical for the generation of broadly neutralizing antibodies, for example for Influenza A virus hemagglutinin (HA) stem-binding antibodies, which require IGHV1-69 alleles containing F55 rather than L55 ^25^, or SARS-CoV-2 RBD binding antibodies that require both R50 and F55 ^29^. Using the 1KGP dataset generated here, we asked if the proportion of individuals with permissive IGHV1-69 alleles for these types of antibody responses differed between the populations. We found that only 5% of Africans lacked F55-containing IGHV1-69 alleles compared to up to 40% of South Asians and 25% of East Asians, which may influence the ability to generate influenza HA stem-directed neutralizing antibodies (**Figure 6F**). We found that the 1KGP African individuals had the highest frequency of IGHV1-69 alleles containing both R50 and F55 (**Figure 6G**). Furthermore, in a recent study, we demonstrated that IGHV2-70 alleles with a CDR2-encoding R52 were non-permissive for elicitation of influenza neutralizing antibodies of the LPAF-a class ^26^. We identified 11 variants with unique coding sequences for IGHV2-70 (**Figure S6B** and **D**) and 10 for IGHV2-70D (**Figure S6C** and **E)** with either R52 or L52. Analysis of IGHV allele frequencies in the 1KGP set revealed population differences that may influence elicitation of LPAF-a class responses (**Figure 6H**).

### A *de novo* meiotic recombination event created a novel IGH haplotype

The IGH locus is distinguished from most other genomic regions by the presence of both high degrees of allelic and structural variation, and the presence of multiple pseudogenes. These features are consistent with a model that involves high levels of meiotic recombination events, resulting in gene duplications and deletions, allowing more frequent generation of novel haplotypes compared to most other genomic regions. Haplotypic analysis of a family group consisting of two parents and two children (**Figure 7A**) provided an example of *de novo* haplotypic generation. The IGHV genotypes of the two children were consistent with all alleles being inherited from either of the parents (**Figure 7B**). All family members were heterozygous for IGHJ6*02 and *03, which enabled their use as inferred haplotype anchors and mapping of the IGHV alleles to each chromosome (**Figure S7A-E**). Critically, the father had two IGHV haplotypes, one of which contained a large 13 gene deletion between IGHV3-7 and IGHV3-30, in addition to a shorter structural deletion that included the 5 genes between IGHV3-30 and IGHV4-34 (**Figure 7C**). This haplotype also contained a rare IGHV1-69 allelic variant, IGHV1-69*04_S8897, containing an isoleucine at position 55. Our analysis showed that the deleted IGHV haplotype, including the IGHV1-69*04_S8897 allele, was inherited by the daughter, in addition to one of the haplotypes from the mother (**Figure 7C**). In contrast, the son inherited a haplotype from the mother in addition to an additional haplotype that differed from either of the 4 parental haplotypes. Upon closer examination of this additional haplotype, it was apparent that it contained the family-specific IGHV1-69*04_S8897 allele. Examination of the overall allelic content was consistent with a meiotic recombination event between the two haplotypes present in the father, juxtaposing the chromosomal region between the genes IGHV3-53 and IGHV1-58 (**Figure 7D**). Of note, this *de novo* recombination event resulted in the ‘repair’ of the 13 IGHV gene deletion and added 3 of the 5 genes that were absent in the shorter deletion, demonstrating an ongoing genomic process for continued IGH locus diversification captured in “real time”.

**Figure 7.**
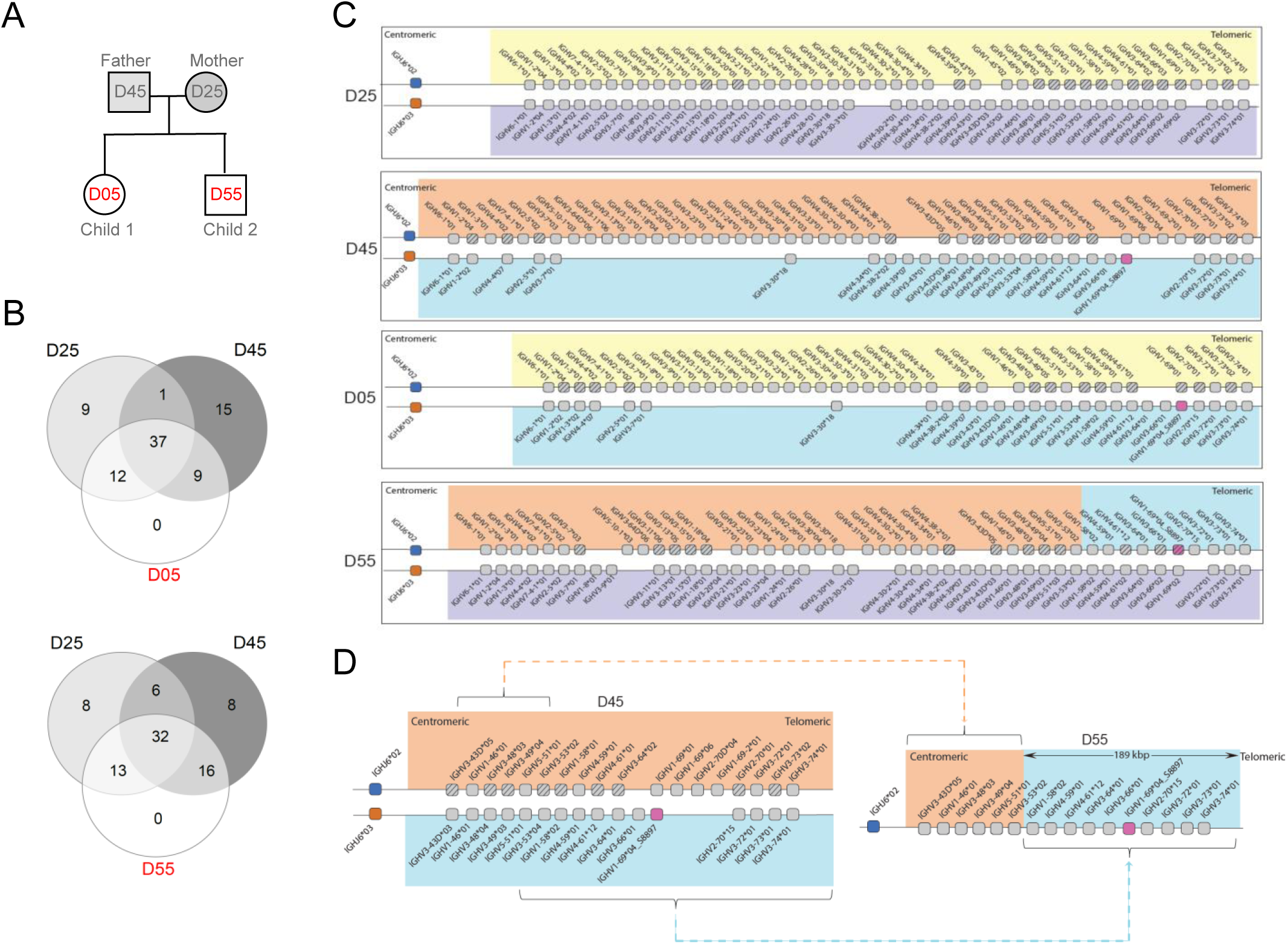
Identification of *de novo* haplotype recombination. **(A)** Family tree of two parents (D45 and D25) and two children (D05 and D55). **(B)** IGHV allele content of parents and children. **(C)** Schematic of the inferred IGHV haplotypes based on heterozygous IGHJ6*02/*03 anchoring for each of the four individuals. IGHV1-69*04_S8897, indicated with a red shaded box, is unique to this family within this study. **(D)** Schematic illustrating the recombination in the two haplotypes present in parent 2 between IGHV3-53 and IGHV1-58, resulting in a new IGHJ6*02 associated haplotype in child 2 that contains the entire gene content telomeric of IGHV3-53 breakpoint, including the IGHV1-69*04_S8897 allele.

## Discussion

The extent of adaptive immune gene heterogeneity at the individual and population level has begun to be uncovered in recent years ^3,4,35,36,60,61^, along with the realization that germline variation can profoundly affect paratope-epitope specificity ^25,29–31,62^. The recent SARS-CoV-2 pandemic provided a sobering lesson that emergent diseases can rapidly impinge on all global populations, that a deeper knowledge of antibody responses is a critical factor in how we address this problem, and critically, that a comprehensive understanding of the existing functional variation of these genes is currently absent. We addressed this need through the development of a highly accurate and high throughput sequencing approach, ImmuneDiscover, designed to identify all IGHV, IGHD and IGHJ germline variants present within large cohorts of individuals. Applying this to 2486 individuals of the 1KGP samples, a population set highly representative of global population variation, enabled the determination of allelic frequencies across 25 different global populations. The assembly of this data set provides an important research resource, KIARVA, that has numerous implications for future immunological studies.

The best-known example where IGHV allelic variation was shown to influence epitope recognition is influenza A virus HA central stem-targeting broadly neutralizing antibodies that use IGHV1-69, which require alleles that encode a phenylalanine (F) in position 55 of the HCDR2 ^25^. In the accompanying paper, we describe a further example of allelic biases involving a class of neutralizing antibodies targeting the HA head that require IGHV2-70-alleles with a leucine (L) in the HCDR2 ^26^. Studies performed during the SARS-CoV-2 pandemic provided additional examples of how germline-encoded variations in IGHV genes influence epitope recognition. Through monoclonal antibody isolation and germline allele reversions, we reported that a class of receptor binding domain (RBD)-targeting neutralizing antibodies required the presence of both R50 and F55 in IGHV1-69*20 ^29^. Furthermore, Yuan et al. reported that IGHV2-5*02-using neutralizing antibodies against RBD binding, required alleles with D54, while N54-encoding alleles were non-permissive for this binding mode ^31^.

The IGHV allelic frequencies provided in KIARVA provides important information relevant to ongoing germline-targeting vaccine trials for HIV. When the goal is to elicit VRC01 class antibodies, IGHV1-2 alleles that encode W50 (IGHV1-2*02 and IGHV1-2*04) are permissive, while alleles that encode R50 (IGHV1-2*05 and IGHV1-2*06) are non-permissive. Four independent vaccine trials held in the US, Europe and Southern Africa, in which personalized IG genotyping was performed, have confirmed that individuals who lack permissive IGHV1-2 alleles were non-responsive to the vaccines ^32,56–58^. The current analysis of the 1KGP set demonstrated that IGHV1-2*05 exhibits strong population bias, with over 20% of both South Asian and East Asian populations lacking a VRC01 permissive IGHV1-2 allele. Biases for specific IGHV genes have been described in the response to several additional pathogens, including IGHV5-51, IGHV3-13 and IGHV3-23 for *Human parainfluenza virus 3 (HPIV3),* Ebola virus and Zika virus surface glycoproteins, respectively ^63–65^.

It is important to note that many IG alleles are not population-specific, as the populations themselves are heterogeneous in terms of gene and allele content. However, the frequency of specific allelic variants differs widely between populations studied, which may have implications for global vaccination strategies. For example, antibody responses that require IGHV3-33, IGHV3-9, IGHV1-69-2, IGHV4-38-2, IGHV4-31, or IGHD genes located within the IGHD3-3 to IGHD2-8 deletion, will result in a lower response in populations with a higher frequency of these structural deletions. Overall, two major features govern the ability to recognize target epitopes, the HCDR3, in which IGHD gene usage is of prime importance and the HCDR1 and HCDR2, which are germline-encoded by each IGHV gene. The current study demonstrates that germline-encoded variations that differ in frequency between human population groups affect both HCDR3 and non-HCDR3-mediated binding modes. The IGHD3-3-IGHD2-8 multigene deletion while found in all populations, is present at a frequency of up to 30% in East and South Asian populations. The deleted haplotype results in the absence of several highly utilized IGHD genes in homozygous individuals, including IGHD3-3, IGHD4-4 and IGHD6-6. Importantly, each of these genes contains sequences that are not present in other expressed IGHD genes, suggesting that the HCDR3 repertoires of deleted versus non-deleted individuals are functionally different in naïve repertoires.

Natural IgM antibodies represent another level of germline-associated functionality where IG genes that are frequently used in naïve repertoires, such as IGHV1-69, IGHV3-21 and IGHV1-2, form classes of unmutated ‘public’ antibodies. Natural antibodies have been found to bind common bacterial, viral or parasite derived targets without the need for affinity maturation, thus enacting a more innate-like function ^66,67^. They also play a role in the clearance of apoptotic cells that express self-antigens and through this function have been linked to autoimmunity ^68^. The impact of IG polymorphisms on the function of natural antibodies remains unknown.

The care taken to ensure the KIARVA dataset contains variation present in the larger human population, is designed avoid the presence of erroneous sequences, the inclusion of which within a human reference set may negatively impact various analyses. We suggest that many previous efforts to understand basic features of the human antibody repertoire that used human reference sets that were inadequately curated to avoid erroneous content may be improved upon based on the highly curated information provided in KIARVA, such as a recent analysis of codon stability coefficient scores of IGHV germline genes ^69^.

Comparison of the IG genes between different vertebrate species indicate the IG loci have undergone evolutionary changes specific to individual species ^70,71^. This variation may be driven by multiple factors, both neutral due to founder effects within an original heterogenous population, or adaptive, in response to endemic pathogens. Our understanding of how pathogens have interacted with, and ultimately shaped our germline gene repertoire, requires knowledge about IG gene and allele frequencies in different populations and, a greater challenge, how these have changed over time. The current study provides an estimate of current IG frequencies in the major human populations and facilitated the collation of a comprehensive set of specific allele-associated SNP variants. The vast majority of the SNP variants found in this study are absent from current SNP arrays commonly utilized in GWAS analyses ^14^, a factor that may have limited our ability to detect IGH related disease associations to date. A comprehensive set of IGH SNPs will facilitate the development of an adaptive immune SNP collection for GWAS studies. Importantly, the utilization of this SNP analysis with archeological DNA samples ^72^ may provide a means to identify gene frequency changes in populations that have undergone migration-associated population changes. It is of note that many of the novel IGHV variants we identified in a recent analysis of South African individuals ^73^ were also present in the West and East African populations in the 1KGP set, presumably due to shared Bantu heritage across sub-Saharan populations ^74^. Comparison of gene frequencies from different historical timepoints may uncover adaptive immune adaptive selection due to historical pandemics, for example following the European medieval bubonic plague, the spread of malaria strains across the world, or the post-Columbian American depopulation period following European disease transmission to the Americas ^75–77^.

The level of human IGH genetic diversity may be intimately related to the chromosomal positioning of the gene at the telomeric end of chromosome 14. The IGHV locus is present in a telomeric location in multiple species, including the non-human primates, cat, dog, mice and rats, suggesting that this genomic positioning provides a structural advantage. One possible mechanism involved may be the ability to generate additional variation in the locus through an increased rate of recombination. Recent analysis using whole genome sequencing of multiple family members revealed a 50-fold difference in meiotic recombination rate within 2 Mb of telomeres in males compared to females ^78^. The family analysis described in the current study illustrates this point, with the meiotic crossover occurring in the male line, and critically, provides a potential explanation for how populations maintain genetic diversity. The crossover event between the deleted and a non-deleted haplotype in the male parent (D45) described here resulted in the recombined haplotype observed in child D55. This enabled the retention of multiple distal gene variants that were originally present in the deleted haplotype in D45. These allelic variants are present on a *de novo* haplotype that retains the previously deleted segment of 11 IGHV genes. The ability to generate *de novo* IGHV non-deleted haplotypes through the recombination of chromosomes containing structural deletions is dependent on population heterozygosity.

While IGH structural deletions are present at different frequencies in the larger superpopulation groups, variation also exists *within* populations, suggesting the possibility that many variants exist within a background of balancing selection. How differences in IG gene content between human populations rather than prior immunological exposure impact the ability to respond to a pathogen or auto-antigen remains largely undefined. Population reduction in the Americas following the European migration in the fifteenth century shows that pathogen transfer to an insufficiently immunologically experienced population can be significant. Likewise, we know that archaic human population groups declined at a time when modern populations migrated from Africa to the Middle East, Europe and Asia, encountering Neanderthal and Denisovans in so doing. Interestingly the highest degree of IGHV deletions, as determined by the gene number found in the later Neanderthal samples, with the lowest found in the Vindija case, the most recently surviving Neanderthal individual for which a high coverage assembly is available ^46^, is indicative of restriction in population numbers of the later Neanderthals, with concomitant loss of genetic diversity. Whether this loss of adaptive immune genes in Neanderthals resulted in a less efficient adaptive immune response compared to the more genetically diverse out-of-Africa human population they encountered is, at present, an open question.

In conclusion, we present multiple critical findings that will guide future studies of the role of immunogenetic variation for disease susceptibility and accelerate initiatives in precision medicine, as well as provide insight on the evolution of the human IGH locus. Our results demonstrate that each person has their own set of IG alleles, shaping their BCR repertoire. As we have indicated here, disease specificities are in the process of being defined for the range of IG variants, resulting, ultimately, in the possibility to identify individual immunological vulnerabilities based on an IG genotype. Finally, the ImmuneDiscover technique enables population-scale IG genotyping with unprecedented resolution, opening new avenues for investigating how antibody gene heritability shapes immune responses, disease susceptibility, and therapeutic outcomes across populations

## Limitations of study

The current study is restricted to the analysis of functional IG genes in 25 population groups from the 1KGP cohort and may miss variation that is present in unsampled populations. This includes both the superpopulation (continental) level, such as African and South American populations and in the limited or absent sampling of regions that contain large numbers of individuals, such as South-East Asia. To limit the possibility of inclusion of SHM derived false positives, the study database includes only those 1KGP derived alleles that were found in at least two individuals, with the result that rare variants found only in a single individual are missing from KIARVA. Future analysis with larger sample sizes will enable such variants to be corroborated. The use of the lymphoblastoid cell line-based 1KGP set restricts our ability to identify all variants of IGHV2-70/IGHV2-70D, as well as the IGHD and IGHJ genes. While the more frequent variants in these genes were identifiable using the filters for ImmuneDiscover analysis used here, additional variation may be present.

### Supporting information

Supplemental Figure 1

Supplemental Figure 2

Supplemental Figure 3

Supplemental Figure 4

Supplemental Figure 5

Supplemental Figure 6

Supplemental Figure 7

## Acknowledgements

We thank Anton Sendel for sample collections, the study participants for providing samples and Xaquin Castro Dopico for helpful comments on the manuscript. Funding for this work was provided by a Distinguished Professor grant from the Swedish Research Council (grant 2017-00968), an ERC Advanced grant (grant 78816) and a Wallenberg Scholars grant (KAW 2023.0248) from Knut och Alice Wallenbergs Stiftelse to G.B.K.H. We are grateful to Fondation Dormeur, Vaduz for support of equipment. The KIARVA resource was developed by the Data Science Node in Precision Medicine and Diagnostics, which is part of the SciLifeLab & Wallenberg Data Driven Life Science Program, funded by Knut and Alice Wallenberg Foundation (grant 2020.0239).

## Author contributions

Conceptualization: M. Corcoran and G.B.K.H.; Methodology: M.Corcoran developed the ImmuneDiscover library production technique and M.K. wrote the ImmuneDiscover analysis script and developed IgSNPer. M. Chernyshev compiled data for the KIARVA resource. Investigation, analysis and visualization: M. Corcoran, S.N., M.K., M. Chernyshev and G.B.K.H.; Sample resources, A.F. and C.S.; Funding acquisition, G.B.K.H.; Supervision, M. Corcoran and G.B.K.H.; Writing original draft: M. Corcoran and G.B.K.H. All authors reviewed, edited and approved the submitted manuscript.

## Methods

### Experimental subject details

### Human subjects and ethical statement

The KI cohort consisted of participants who self-identified as having ancestry of either of 4 superpopulations (EUR, AFR, EAS or SAS). The inclusion of study participants was approved by the Swedish Ethical Review Authority: decisions 2016/1053-31, 2017/852-32, 2006/893-31/4, 2011/222-31/1, 2013/549-32, 2013/550-32, 2018/2354-32, 2019-01014). All samples were collected with written informed consent, pseudonymized and handled according to regulations.

## Method details

### RNA and DNA isolation

Blood samples were collected in BD Vacutainer K2 EDTA tubes and PBMCs were isolated using Ficoll-paque^™^ (GE healthcare life sciences). From the African cases we obtained frozen PBMCs for RNA extraction using the RNeasy kit (Qiagen). Genomic DNA was extracted using Gentra Puregene Buccal Cell Kit (Qiagen).

### 5’ Multiplex PCR for IgM library analysis

IgM libraries were prepared as previously described ^38^. Briefly, 200ng of total RNA was used for reverse transcribed into cDNA using an IgM chain-specific primer that contains Read2 sequence and a UMI using the Sensiscript RT kit (Qiagen). We designed two independent primer sets for each of the V gene that bind to either the 5’ forward leader (leader primer set) or the 5’ UTR (Upstream primer set) to produce two independent libraries containing full length VDJ constructs. The cDNA reaction was purified (PCR purification Kit, Qiagen) and PCR amplified using the Read2U reverse primer and either the 5’ Leader or Upstream primer mix. The PCR product of approximately 500 bp was gel purified using the Gel purification kit (Qiagen).

### NGS library preparation and sequencing

Five nanograms of the gel-purified template was used for the index PCR reaction. The forward index primer (P5_R1) and reverse index primer (P7_R2_1-27) was added to the 10-cycle PCR reaction along with KAPA HiFi Hotstart readymix system (Kapa Biosystems). The libraries were purified first using PCR purification kit (Qiagen) followed by magnetic bead purification (AMPure XP magnetic beads, BD). The libraries were quantified and sequenced using Illumina version 3 (2 x 300 bp) sequencing kit with 13% of 12 pM PhiX172 bacteriophage DNA.

### Computational analysis

The IgDiscover pipeline version v1.0.4 was used for pre-processing of the obtained reads and generation of personalized databases. The AIRR reference database downloaded in November 2023, containing germline alleles of IGHV, IGHD and IGHJ was used as a starting database. We additionally used the *corecount* ^39^ module for accurate genotyping. This module makes use of a set of filtered sequences from the IgDiscover output and derives a set of core V-D-J sequences present within the starting database. Each of these core sequences are searched within the filtered library and the number of sequences carrying the core sequence are counted and filtered to remove false positives to give rise to final genotype.

### High throughput genomic genotype analysis: ImmuneDiscover

The ImmuneDiscover genotype methodology utilizes a customized multiplex set of targeted primers, specific for either the IGHV, IGHJ or IGHD genes. The primers are designed to amplify a product of approximately 450 bp for each target, thereby enabling, for example, the amplification of all IGHV alleles from a single individual in a single well. The primers are designed to target upstream and downstream of non-rearranged germline sequences. The lack of a downstream reverse target sequence means that the multiplex set will only amplify non-rearranged germline sequences and will fail to amplify rearranged VDJ sequences. This enables the technique to be used on DNA templates from multiple sources, including PBMCs that contain many rearranged VDJ sequences. The ImmuneDiscover process has been designed to work using nanogram quantities of target DNA and is performed in a 96 well format, with different individual template DNAs in each well. Each target primer contains a universal forward and a universal reverse overlap sequence to enable subsequent high throughput indexing. ImmuneDiscover-identified alleles were required to pass donor count thresholds to be included in KIARVA. Each allele had to be present either in one of the KI donors and/or two or more of the 1KGP donors. During our review of ImmuneDiscover inferred alleles from 1KGP donors, we found an unusually high number of IGHV2-70 and IGHV1-69/1-69D alleles. These alleles appeared to be the result of off-target SHM resulting from the genomic proximity of the two genes, requiring more stringent donor count thresholds for these genes. Specifically, we required alleles to be present in at least 10 donors for IGHV2-70 and at least 3 donors for IGHV1-69/1-69D.

### Multiplex indexing

The ImmuneDiscover process utilizes a novel indexing procedure. The forward and reverse universal overlaps enable the addition of forward and reverse well specific indices, each of which is specific for one well of the 96 well plate. In addition, a plate specific index is added at this stage, resulting in each library having two well specific indices and one plate specific index. This allows the multiplexing of libraries from multiple plates during an NGS sequence run. For the indexing procedure, 5µl of the library PCR is purified using Ampure magnetic beads (AMPure XP magnetic beads, BD) and resuspended in 40 µl of H2O to remove the library PCR primers. 4 µl of this is added to the index PCR step, which is allowed to proceed for 12 cycles, thereby adding the indices to each library. The libraries from a single plate are pooled and gen purified, quantified using Qubit fluorometer and sequenced using the MiSeq Illumina V3 2 x 300 cycle kit along with 13% 12 pM PhiX172 DNA.

### ImmuneDiscover computational pipeline

The ImmuneDiscover program consists of a set of distinct subcommands which are part of the pipeline for genotyping and discovery of novel alleles from massively parallel sequencing runs. The demultiplexing process is the initial step in the ImmuneDiscover pipeline. This key step segregates the input FASTQ file into appropriate cases based on the provided indexing information. The process requires a compressed FASTQ file (fastq.gz) and a tab-separated (TSV) file containing mandatory columns for forward_index, reverse_index, and case. The demultiplexing command extracts reads with the indexing barcodes provided in the indices file, matching the forward index at the 5’ end and the reverse index at the 3’ end. The output is a compressed TSV file containing columns for well, case, read name, and genomic sequence. This output serves as the foundation for subsequent analysis steps. The demultiplexing process can be fine-tuned using various parameters. Users can specify a minimum read length and optionally split the output into separate FASTQ files per case. These options provide flexibility in handling diverse experimental setups and data structures.

Following demultiplexing, ImmuneDiscover employs an exact match search to identify known alleles within the sequencing data. This step requires demultiplexed TSV and a FASTA file containing a database of known alleles. The exact match search algorithm compares each read against the provided database, identifying identity matches. The process distinguishes between "collapsed records" (matches excluding flanking regions) and "full records" (matches including flanking regions). This distinction allows for a more nuanced analysis of allelic variants outside of the query sequence (i.e. recombination signal sites). The exact match search output is a TSV file containing detailed information about each identified allele. This includes the well and case identifiers, database allele name, sequence, counts (both full and collapsed), gene association, and various frequency and ratio calculations. Additional columns may be included based on the specified gene option, providing information on sequence elements such as heptamer, spacer, and nonamer regions. The exact match search process can be customized using several parameters. Users can set minimum allelic ratios and specify reference genes for ratio computations.

The final major component of ImmuneDiscover is the BLAST-based novel allele discovery process. This step is designed to identify potential novel alleles that are not present in the existing database. The process involves running BLAST ^79^ assignments on the sequences, followed by a series of filtering and analysis steps. For V genes, the process performs BLAST assignment, collapses identical sequences, and applies filters to identify novel alleles. For D genes, the process is more complex due to their short lengths. It first extends the sequence by a specified length, uses the most common occurrence of this extended sequence as a BLAST query, and then re-aligns the results to the original database using semi-local alignment. This approach allows for the restoration of the original sequence length and accurate determination of mismatches in the original non-extended gene. The BLAST process output is a compressed TSV file containing detailed information about potential novel alleles. This includes well and case identifiers, BLAST-assigned allele names, mismatch counts, read counts, gene associations, frequencies, and alignment information. The output also includes a new allele name, which may be tagged as "Novel" if the allele is determined to be previously unidentified. The BLAST-based discovery process can be fine-tuned using various parameters. Users can specify minimum frequency thresholds, minimum count, maximum allowed distances for alleles, and minimum read lengths. For D genes, users can also specify the length of sequence extension. These parameters can be used to adapt the pipeline to the specific requirements of experimental setup. As part of the analysis pipeline, ImmuneDiscover offers a separate BWA sub-command ^80^ to align reads to a reference genome. This step is crucial for filtering out pseudo-genes from other chromosomes, thereby enhancing the accuracy of allele identification and novel allele discovery. The ImmuneDiscover software is available from the Karlsson Hedestam software repository.

### IgSNPer analysis

IgSNPer is a pipeline developed to analyze newly discovered alleles by comparing them to specified genomic locations, identifying and quantifying uncommon single nucleotide polymorphisms (SNPs). The tool assigns a score to each query sequence based on the number of uncommon SNPs found, with an uncommon SNP being defined as found below a frequency of <0.0001 in the multiple population dataset. We surmised that SNPs with low population frequencies have a higher chance of being of non-germline origin, particularly if multiple such SNPs are found in the same candidate allele. This is supported by extensive sequencing data from diverse populations aligned to reference genomes, which enables SNP frequency estimation and serves as additional evidence. The analysis requires four types of input files. 1) A mapping file (mapping.json) which maps genomic identifiers to chromosomes and links assemblies (GRCh37/38) to their corresponding VCF and FASTA files. 2) A coordinates file in TSV format lists the gene names, assembly, and coordinates (chr:start-end) manually derived from BLAT mapping of allele variants to the GRCh37/38 assemblies for accurate reference assembly positioning. 3) Query sequences are provided in FASTA format, where identifiers must match gene names in the coordinates file. 4) Reference data that includes GRCh37 and GRCh38 genome assemblies along with dbSNP VCF files for both assemblies.

The pipeline itself consists of multiple stages. During data preparation, sequences are converted to a tabular format while gene names and locations are extracted to create gene-specific groups (_group.tsv) and generate query FASTA files (_query.fa). Reference processing involves extracting reference sequences based on coordinates (_reference.fa), performing initial BWA alignment for sequence positioning to generate SAM files, and extracting relevant dbSNP variants (_variants.tsv). The alignment and variant detection stage employs semi-global alignment using the Smith-Waterman algorithm with gap penalties of open=-50 and extend=-1, utilizing the EDNAFULL scoring matrix. Query alignments are compared to reference sequences to identify variants. For variant classification, each detected variant position undergoes matching against dbSNP variants from VCF files. When a unique match is found, frequencies are checked in population databases from the VCF file, with variants classified as common if frequency ≥ 0.0001 and uncommon if frequency < 0.0001 in the combined datasets. Variants are assigned rsUNK identifiers when no dbSNP match is found or when multiple rs identifiers are reported at a position without a unique variant nucleotide match to neither of reported in VCF. Population frequency analysis follows a specific hierarchy of datasets checked in the following order: dbGaP_PopFreq (>1 million subjects), ExAC (60,706 individuals), TOPMED (158,000 individuals), TOMMO (8,380 Japanese individuals), KOREAN (1,465 individuals) and, GoESP (6,503 individuals). The reported rs number originates from the first dataset where a SNP matching the identified variant in the alignment between query and reference was found. The order is to prioritize larger, more reliable datasets.

Several special handling procedures are implemented within the pipeline. For indels, long gap stretches are allowed only within sequences, while edge gaps are treated as truncations and counted as independent uncommon SNPs reducing score for incomplete alleles. The analysis excludes 1KGP containing datasets to ensure independent evidence for frequency determination. The pipeline generates a TSV file containing the allele name, number of uncommon variants, list of uncommon variants with rsID/rsUNK identifiers and variant positions (variant:aln_pos,genome_pos), number of common variants, and list of common variants with rsID identifiers and positions (variant:aln_pos,genome_pos). The program can analyze all human variants presented in KIARVA except for alleles of IGHV4-38-2, as this gene is not mapped to coordinates within the primary GRCh37/38 assemblies and is therefore missing the corresponding gene specific SNP data.

### D-J clonality analysis

Additional analysis examines polyclonal immune receptor diversity across different human populations using custom DNA probes. The probes were prepared manually using BLAT (BLAST-Like Alignment Tool) and the D and J gene databases to select each probe such that it would match at least one D or J gene segment from the immunoglobulin heavy chain locus. This targeted approach allows for specific capture and identification of D-J recombination events in genomic data from diverse population groups represented in the 1KGP. The workflow involves analyzing sequencing reads that match these custom probes, followed by identifying sequences matching D and J probes on the same read. For each donor from each population, samples were categorized based on the presence of multiple distinct D-J recombination patterns. The polyclonal metric was computed by examining all unique D-J sequences within each sample and determining whether more than one unique recombination pattern was present, indicating polyclonality. The frequency of polyclonal samples was computed for each population as evidence of diversity.

### Additional software

Data from multiple IgDiscover runs were collected using the built-in python3.0 script (0.24.2). All the plots were made using Rstudio v4.4.1 with the following packages: cowplot v1.1.3, ComplexHeatmap v2.20.0, Biostrings v2.72.1, seqinr v4.2.36, ggtree v3.12.0, ggplot2 v3.4.4, patchwork v1.2.0, gridExtra v2.3, data.table v1.16.0, tidyr v1.3.1, dplyr v1.1.4 and BioVenn v1.1.3. The circular dendrogram for D alleles was made using Aliview v1.27 and figtree v1.4.4. The alignments were made using BioEdit sequence alignment editor v7.0.5.3. The Venn diagram for the five superpopulations was made using an online tool: https://bioinformatics.psb.ugent.be/webtools/Venn/.

