## Supplemental Figure 1 for "Human IGH germline gene diversity and allele frequencies in 2486 individuals from 25 global populations delineated by ultra-high throughput genotyping"

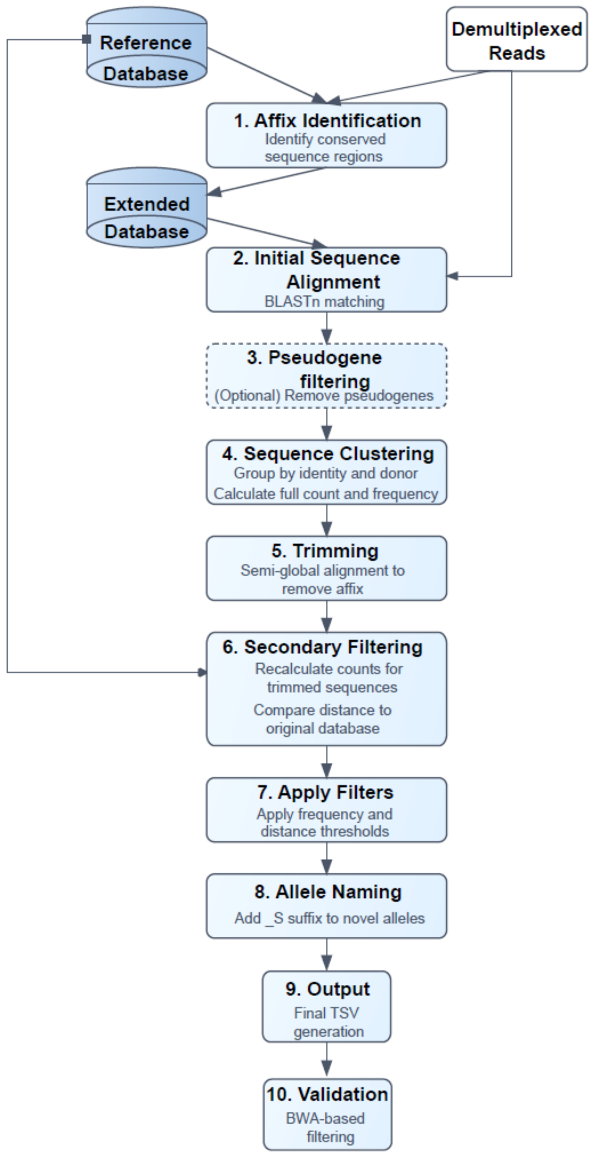


**Figure S1. ImmuneDiscover germline allele discovery computational pipeline, related to Figure 1.**

The ImmuneDiscover computational pipeline performs demultiplexing of the well- and plate-specific amplicons to separate out the individual libraries. The identification of novel germline gene variants is performed using the BLAST module, where demultiplexed sequences are formatted into a FASTA file and aligned to a reference database using NCBI BLAST. Reference databases comprise one with variants of functional genes and one with variants of pseudogenes. Prior to initial assignment, reference sequences are extended by forming an affix on both 5' and 3' ends to capture additional variability using the most abundant affix sequences to create an "Extended database" (**step 1**). After assignments to the Extended database, filtering is applied to separate the best matches to functional genes and identify sequences derived from pseudogenes. Filtering is based on alignment quality, including edge distance (to ensure that variant nucleotides are not at the extreme end of a sequence read), subject coverage, and mismatch count (**step 2**). Additionally, pseudogene filtering can be applied (**step 3**). The filtered sequences are clustered by well, case, target gene, and sequence identity, followed by counts and frequencies calculation (**step 4**). The clustered sequences are trimmed to ensure that the appropriate 3' and 5' affix sequences are separated from each gene sequence (**step 5**); the trimmed sequences are used to recalculate the relative frequency of identified gene assigned clusters, and the sequence distance between each of those clusters and the closest known reference sequence (**step 6**). An allelic ratio filtering is applied to remove clusters that are present below an expected cutoff frequency (**step 7**); the remaining clustered sequences are named as either identical to a known reference sequence, or, in the case of novel variants, it is given a name based on the closest reference sequence with an additional _S suffix (**step 8**); the sequences are finally saved to TSV output (**step 9**) and filtered again using a genomic reference sequence assignment process to ensure that they are derived from the IGH locus rather than a duplicated pseudogenic region on a different chromosome (**step 10**).
