## Supplemental Figure 2 for "Human IGH germline gene diversity and allele frequencies in 2486 individuals from 25 global populations delineated by ultra-high throughput genotyping"

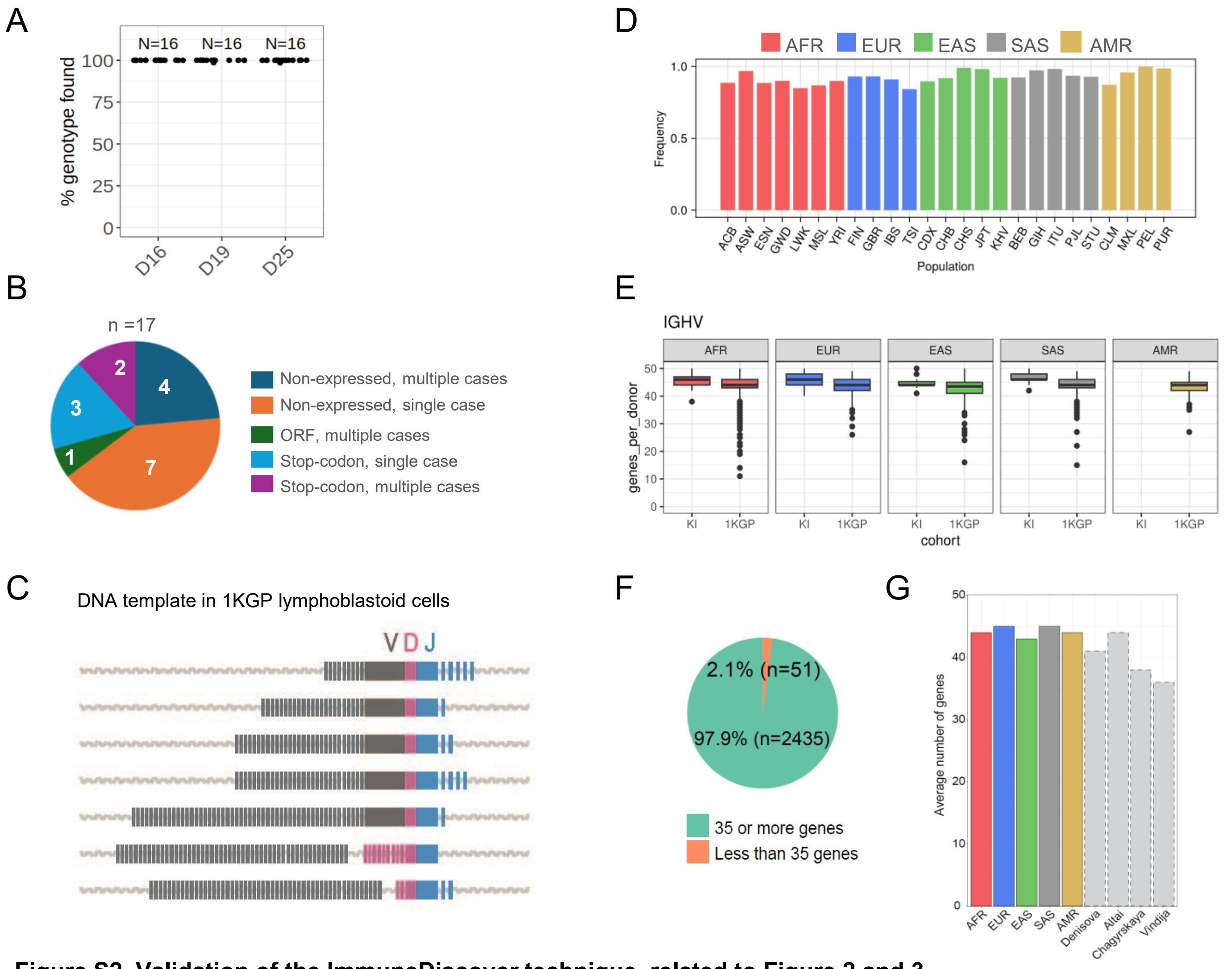


**Figure S2. Validation of the ImmuneDiscover technique, related to Figure 2 and 3.**

**A.** Analysis of reproducibility of ImmuneDiscover genotyping of three cases, D16, D19 and D25. In each case, genomic DNA was used as template in 16 separate library amplifications with subsequent ImmuneDiscover genotyping. The chart shows the proportion of each library that contained 100% of the alleles previously identified for each case **B.** Pie chart showing factors underlying the 17 unique non-expressed IGHV alleles identified using genomic genotyping by ImmuneDiscover not identified from the expression-based genotyping. **C.** Schematic illustrating different stages of IGH VDJ recombinations found in lymphoblastoid cell line DNA samples that involve the loss of different parts of the IGHV, IGHD and IGHJ genes in the IGH locus, and DJ recombinations that retain the full IGHV region, while removing some of the IGHD and IGHJ genes. **D.** Frequency of polyclonal samples across human populations from the 1KGP sample set. Bars show the frequency of samples within each population that exhibited multiple distinct D-J immunoglobulin heavy chain recombination patterns, indicating polyclonal immune receptor diversity. Populations are grouped by superpopulation ancestry and x-axis codes represent specific ethnic groups within each superpopulation category. **E.** Numbers of IGHV genes present in the ImmuneDiscover based genotypes of each KI cohort and 1KGP case. **F.** Pie chart illustrating the number of 1KGP cases that produced IGHV genotypes containing alleles from more than 35 genes. **G.** Bar chart illustration of the average number of genes found in the 1KGP genotypes for each of the 5 superpopulation groups in addition to those found in the three high coverage Neanderthal and single Denisovan archaic assemblies.
