## Supplemental Figure 3 for "Human IGH germline gene diversity and allele frequencies in 2486 individuals from 25 global populations delineated by ultra-high throughput genotyping"

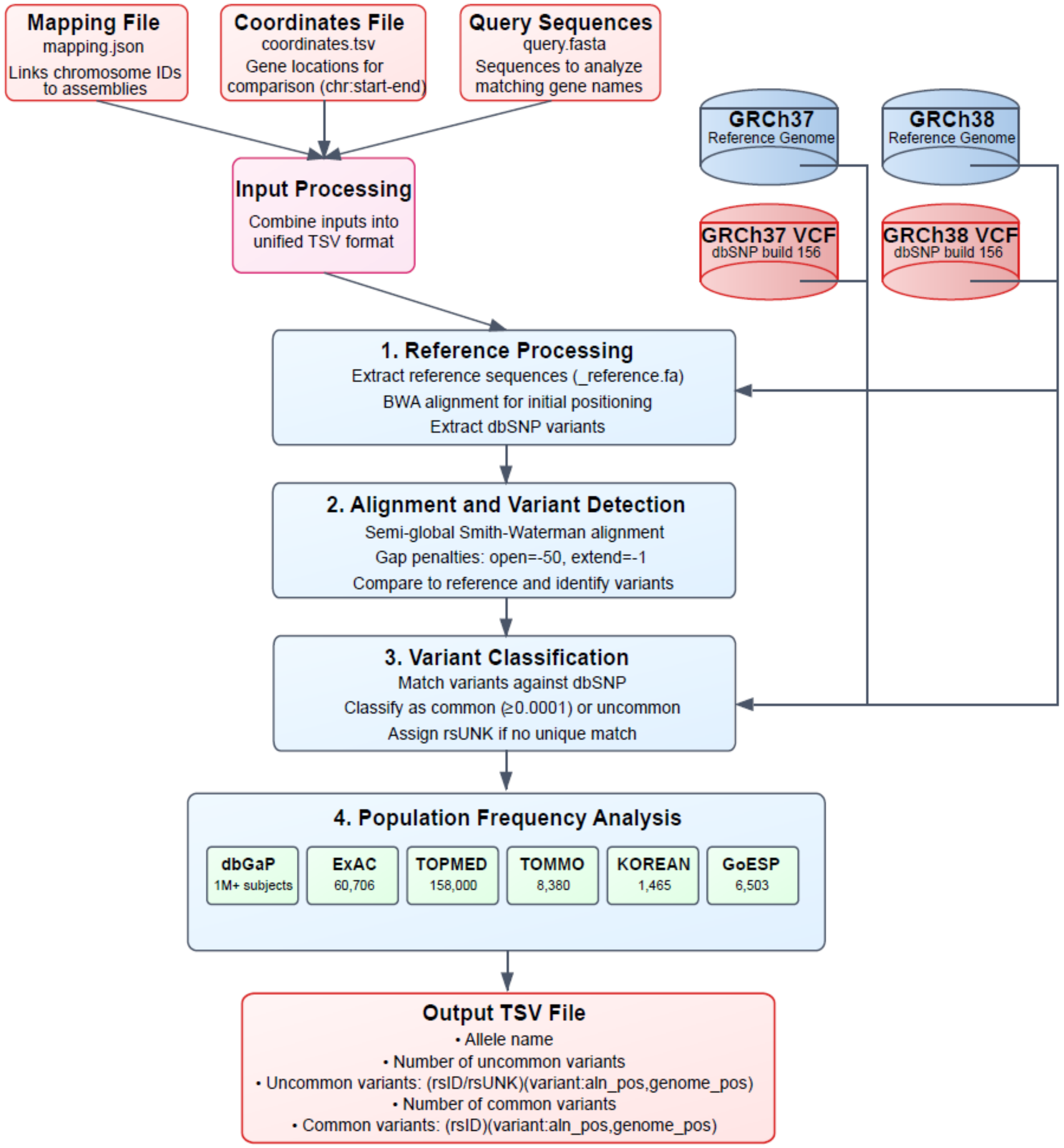


**Figure S3. Schematic of the bioinformatic process involved in IgSNPer analysis, related to Figure 2.** The IgSNPer process utilizes three initial sources of information: a mapping file, containing reference assembly data, a coordinates file, that provides gene specific locations within particular reference assemblies, and a set of query sequences that are the candidate allelic variants being analyzed by the program. The program links the sequences through their ImmuneDiscover assigned gene names, to gene coordinates within specific reference assemblies. The program identifies all variant nucleotides within each candidate allelic sequence by comparing every nucleotide in the sequence to both a reference assembly and to all SNP variants mapped to each position of the gene. The program utilizes a set of prior SNP analyses carried out on collections of multiple individuals to obtain an independent score of SNP variant frequency. IgSNPer utilizes the results of the dbGaP, ExAC, TOPMED, TOMMO, KOREAN and GoESP studies that, in total identified SNP variation in over 1.25 million cases. The IgSNPer program identifies any nucleotide SNP within the coordinates that are absent in the combined SNP population set at a frequency of >0.0001 (defined as uncommon). Each variant nucleotide that is present below the required population frequency increases the IgSNPer uncommon score by 1.
