## Supplemental Figure 4 for "Human IGH germline gene diversity and allele frequencies in 2486 individuals from 25 global populations delineated by ultra-high throughput genotyping"

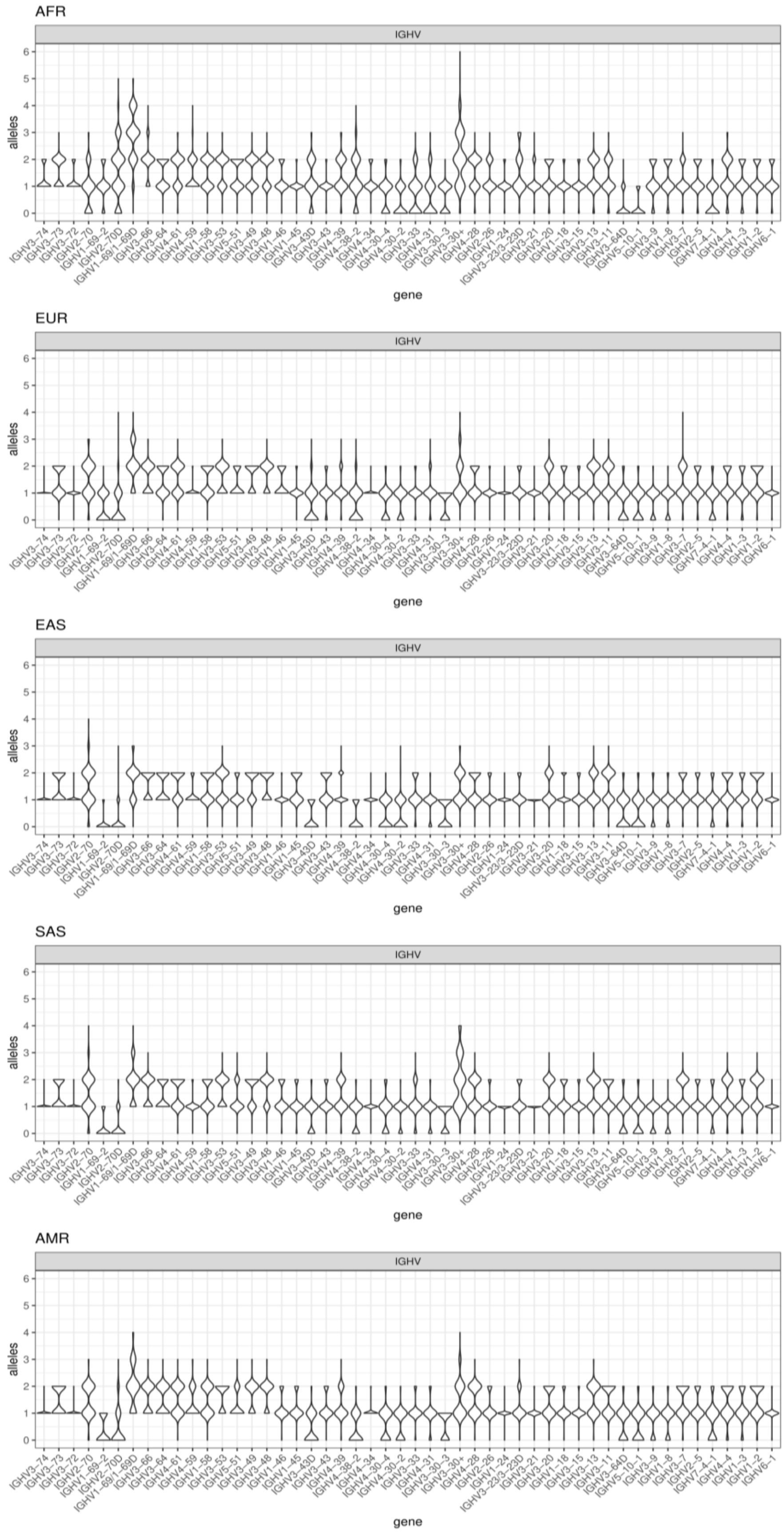


**Figure S4. IGHV copy number in the 1KGP sample set, related to Figure 3.**
Violin plots illustrating the copy number of all functional IGHV genes in each 1KGP superpopulation.
