## Supplemental Figure 5 for "Human IGH germline gene diversity and allele frequencies in 2486 individuals from 25 global populations delineated by ultra-high throughput genotyping"

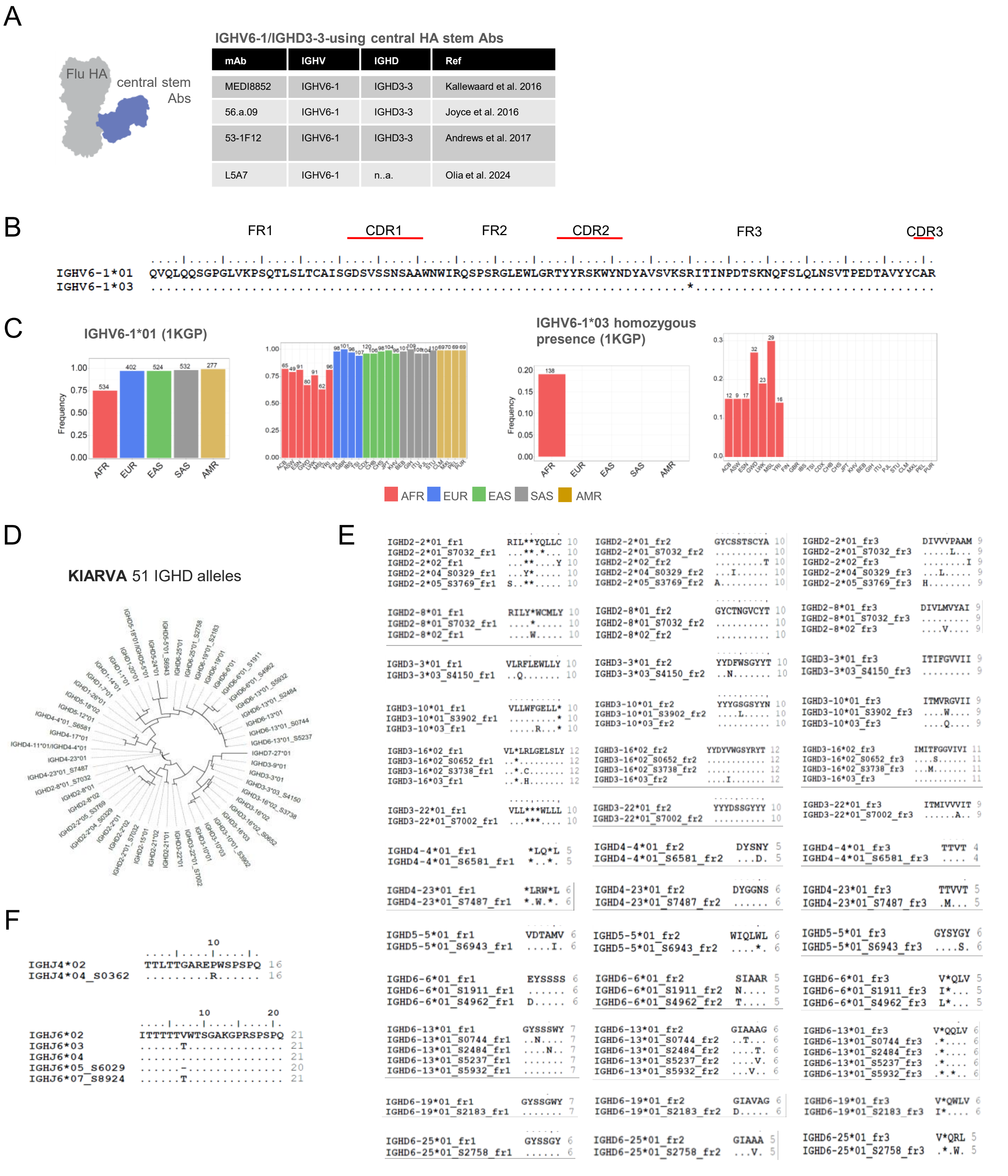


**Figure S5. Alignments and population-frequencies of selected genes, related to Figure 2, 3 and 4.**

**A.** Table of previously described IGHV6-1/IGHD3-3-using influenza HA central stem-binding antibodies. **B.** Amino acid alignment of the frequent IGHV6-1*01 and *03 alleles showing the position of the stop codon in IGHV6-1*03. **C.** Population frequencies of the IGHV6-1*01 and IGHV6-1*03 alleles at the superpopulation level and individual population level. **D.** Dendrogram of all IGHD allelic variants identified in this study. **E.** Amino acid translations of previously unreported IGHD alleles in each reading frame. **F.** Amino acid translations of the previously unreported IGHJ alleles.
