## Supplemental Figure 6 for "Human IGH germline gene diversity and allele frequencies in 2486 individuals from 25 global populations delineated by ultra-high throughput genotyping"

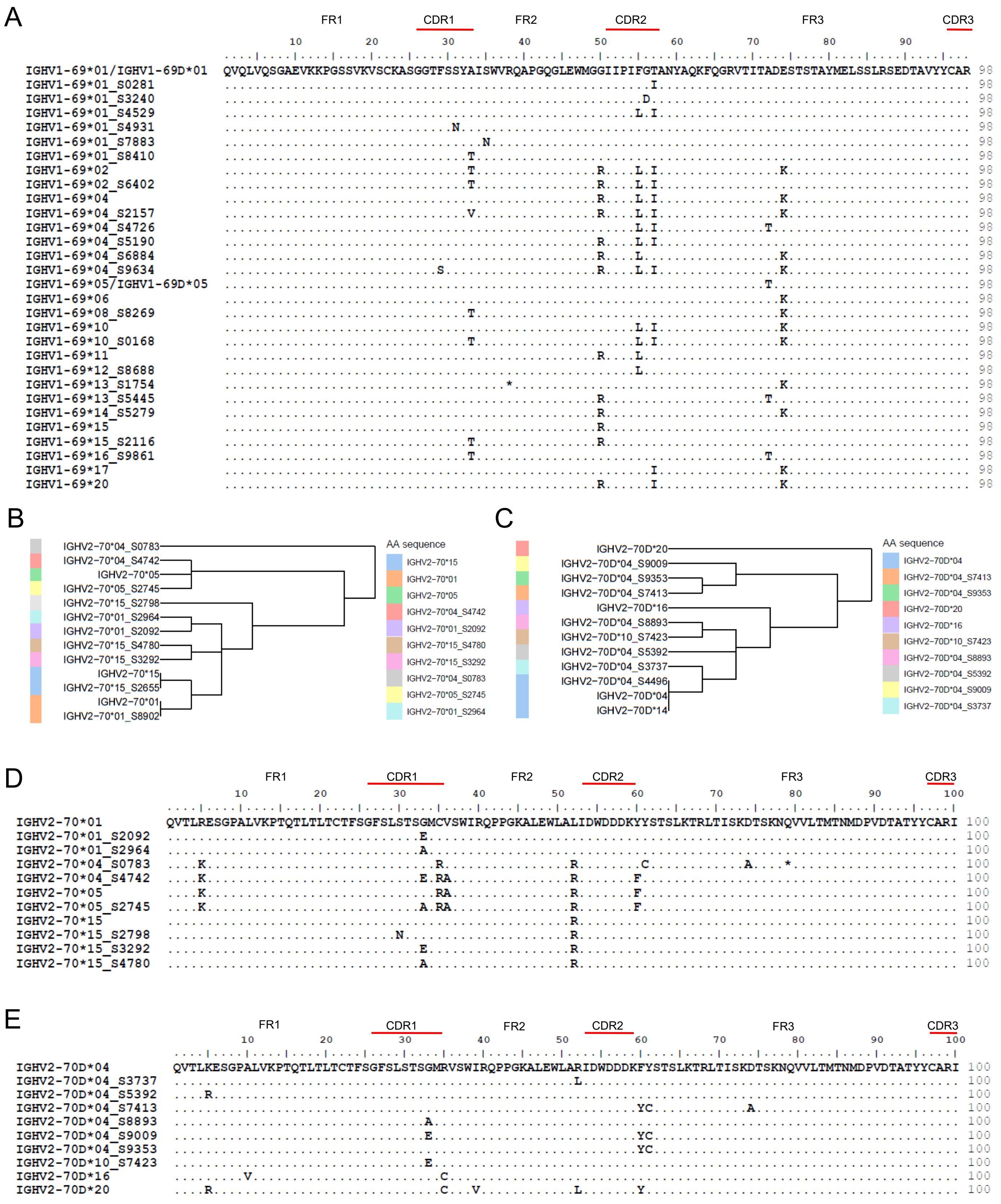


**Figure S6. Amino acid alignments of IGHV1-69, IGHV2-70 and IGHV2-70D alleles with unique coding sequences, related to Figure 6.**

**A.** Amino acid alignment of all IGHV1-69 alleles with unique coding sequences. **B.** Dendrogram of IGHV2-70. germline sequences. **C.** Dendrogram of IGHV2-70D germline sequences. **D.** Alignment of all IGHV2-70 alleles with unique coding sequences. **E.** Alignment of all IGHV2-70D alleles with unique coding sequences.
