## Supplemental Figure 7 for "Human IGH germline gene diversity and allele frequencies in 2486 individuals from 25 global populations delineated by ultra-high throughput genotyping"

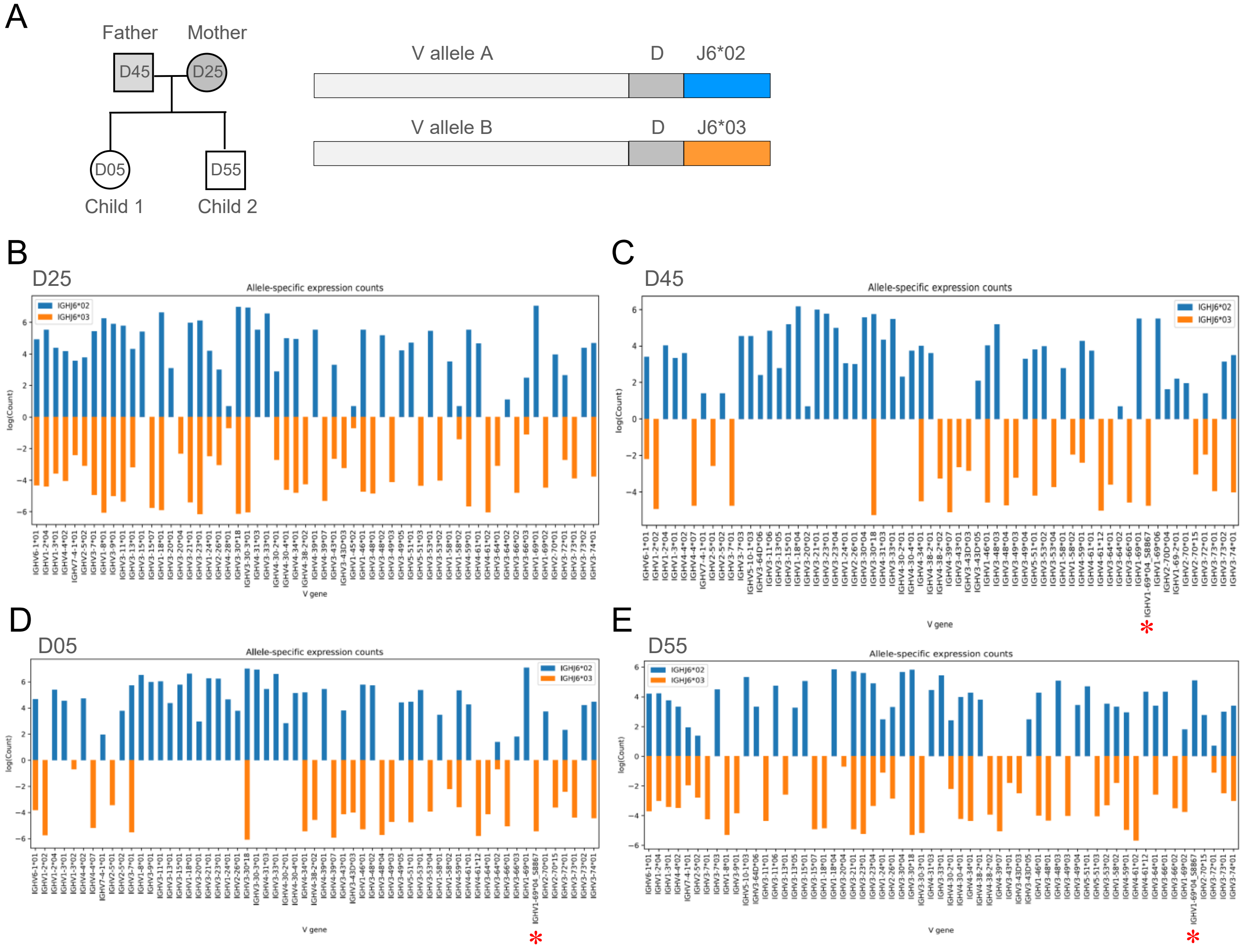


**Figure S7. Inferred haplotype analysis of a family case, related to Figure 7.**
**A.** Schematic of family cases and the use of IGHJ6*02/IGHJ6*03 heterozygosity for J-anchored assignment of IGHV alleles to each chromosome (inferred haplotyping). **B**. Inferred IGHV haplotype of parent D25. **C.** Inferred IGHV haplotype of parent D45. **D.** Inferred IGHV haplotype of child D05. **E.** Inferred IGHV haplotype of child D55. The IGHV1-69*04_S8897 allele is indicated with a red asterix.
